# Shared morphological consequences of global warming in North American migratory birds

**DOI:** 10.1101/610329

**Authors:** Brian C. Weeks, David E. Willard, Aspen A. Ellis, Max L. Witynski, Mary Hennen, Benjamin M. Winger

## Abstract

Increasing temperatures associated with climate change are predicted to cause reductions in body size, a key determinant of animal physiology and ecology. Using a four-decade specimen series of 70,716 individuals of 52 North American migratory bird species, we demonstrate that increasing annual summer temperature over the 40-year period drove consistent reductions in body size across these diverse taxa. Concurrently, wing length – which impacts nearly all aspects of avian ecology and behavior – has consistently increased across taxa. Our findings suggest that warming-induced body size reduction is a general response to climate change, and reveal a similarly consistent shift in an ecologically-important dimension of body shape. We hypothesize that increasing wing length represents a compensatory adaptation to maintain migration as reductions in body size have increased the metabolic cost of flight. An improved understanding of warming-induced morphological changes, and their limits, are important for predicting biotic responses to global change.

## INTRODUCTION

Body size is an essential determinant of animal ecology and life history (Brown 1995; McGill *et al*. 2006), influencing the allometry of physiological (Hudson *et al*. 2013) and morphological (Gould 1966; Outomuro & Johansson 2017) functions, as well as fundamental community ecology interactions (e.g. social hierarchies (Prum 2014), competition, and predator-prey dynamics (Yodzis & Innes 2002)) (McGill *et al*. 2006). Within species, there is evidence that individuals tend to be smaller in the warmer parts of their ranges (an intra-specific derivative of Bergmann’s rule (Bergmann 1847; Rensch 1938; Mayr 1956; Blackburn *et al*. 1999)). This association between warmer temperatures and smaller bodies suggests that anthropogenic climate change may cause intraspecific shifts toward smaller body size in a temporal analog to geographic patterns. However, despite the widespread appreciation of the fundamental importance of body size for ecological and evolutionary processes, the drivers and universality of temperature-body size relationships across space and time remain contested (Riemer *et al*. 2018). Understanding whether rapid body size reductions are occurring in response to increased temperatures is essential to predicting the impacts of climate change on life history, ecosystem dynamics, and the capacity of species to persist in a warming world.

Although the possibility of body size reduction in response to global warming has been suggested for decades (Smith *et al*. 1995; Yom-Tov 2001), empirical support remains mixed (Goodman *et al*. 2012; McCoy 2012; Salewski *et al*. 2014; Teplitsky & Millien 2014; Collins *et al*. 2017a, b; Dubos *et al*. 2018). This uncertainty may be in part due to a scarcity of morphological time series datasets containing sufficiently dense sampling to test the influence of local temporal fluctuations on body size (as opposed to simply associating long-term morphological trends with periods of global warming), and to do so across many co-distributed species that experience similar climatic regimes. Additionally, densely sampled time-series datasets frequently do not have measurements from enough body parts to distinguish changes in body size from changes in body shape that may be driven by alternate selection pressures. Consequently, the influence of warming-driven changes in body size on ecologically-important dimensions of allometry remains largely unknown.

Migratory birds that breed at high latitudes are an important system for understanding the adaptive responses of biota to increasing temperatures, as they are particularly vulnerable to the impacts of climate change. Not only is the most accelerated change occurring at higher latitudes (Soja *et al*. 2007; Contribution to the Fifth Assessment Report of the Intergovernmental Panel on Climate Change 2014), but climate change impacts can vary across the geographically disparate seasonal ranges of migratory species, resulting in complex dynamics such as phenological mismatches between species’ annual cycles and the resources upon which they depend (Charmantier & Gienapp 2014). Migratory birds are under strong selection for high site fidelity, and any perturbation that hinders an efficient return to the breeding grounds is likely to reduce reproductive success (Winger *et al*. 2018). The extreme energetic demands of migration have shaped the morphology of migratory birds for the efficiency necessary to conduct these long-distance flights; therefore, should warming temperatures force body size reductions in migratory birds, concurrent changes in body shape related to the allometry of flight efficiency may be necessary to maintain migratory patterns that have evolved over millennia (Møller *et al*. 2017; Schmaljohann & Both 2017). Although migratory species have garnered significant attention from researchers interested in biotic responses to rapid environmental change, particularly as relates to phenology and geographic range, the extent to which migratory birds are changing size in response to anthropogenic global warming remains uncertain (Van Buskirk *et al*. 2010; Salewski *et al*. 2014; Collins *et al*. 2017a; Dubos *et al*. 2018) and the implications of size change for maintaining physiologically demanding seasonal migrations are unknown.

A persistent challenge in understanding recent morphological changes in migratory birds is the characterization of size and shape (Yom-Tov *et al*. 2006; Salewski *et al*. 2010; Van Buskirk *et al*. 2010). Frequently used indices to assess changes in avian body size through time, such as mass and wing length, are problematic; mass is highly variable for migratory species, given rapid fat gains and losses during migration (Alerstam & Lindström 1990; Morris *et al*. 1996), and wing length is highly correlated with migratory distance (Förschler & Bairlein 2011). Nevertheless, studies on recent body size changes in birds have often represented body size using univariate measures of wing length or mass, making it difficult to identify changes in body size with precision and disentangle them from shifts in shape (e.g. relative wing length) that may be driven by other factors. Wing length is a highly consequential trait in birds that reflects a complex balance of selection pressures from predator avoidance (Witter & Cuthill 1993; Kullberg *et al*. 1996; Swaddle & Lockwood 1998; Martin *et al*. 2018), to flight efficiency (Rayner 1988; Pennycuick 2008), to foraging behavior (Norberg 1979; Fitzpatrick 1985; Miles *et al*. 2002; Ricklefs & Cox 2006). Thus, distinguishing between body size change and shifts in wing length is critical for understanding the ecological consequences of anthropogenically-driven environmental change on migratory birds. This distinction is particularly important as warming temperatures are predicted to reduce body size in birds (Yom-Tov *et al*. 2006; Van Buskirk *et al*. 2010; Gardner *et al*. 2011; Andrew *et al*. 2017, 2018), whereas observed warming-driven changes in migratory phenology, geographic range and habitat (Bowlin & Wikelski 2008; Tingley *et al*. 2009; Förschler & Bairlein 2011; Hahn *et al*. 2016; Møller *et al*. 2017; Socolar *et al*. 2017) have been predicted to select for increases in wing length, potentially resulting in an ecologically-important change in shape (i.e. relative wing length). However, the conflation of wing length and body size has, to-date, largely precluded nuanced analyses of changes in body size and wing allometry (Zink, R. M. and Remsen 1986; Van Buskirk *et al*. 2010).

Here, using a densely-sampled specimen time series of 52 North American migratory bird species, we develop a robust understanding of changes in body size and shape in migratory birds throughout a four-decade period of rapid global change. We take advantage of the ecological diversity of the species studied (see *Ecology and Natural History*, Supporting Information) to test for the presence of consistent morphological change driven by fundamental physiological processes. Specifically, we tested whether increasing temperatures since 1978 have driven reductions in body size. To isolate the impact of temperature on body size, we control for alternate large-scale environmental and climatic variables (precipitation and primary productivity) that could conceivably affect such a diverse set of species. Furthermore, we leverage the multi-decadal and densely-sampled nature of our data to test the influence not only of long-term trends in temperature but also of short-term fluctuations, and in doing so test causal factors of body size change. The multidimensional nature of our mensural data further allowed us to also test how relative wing has changed over the same time period alongside body size. Species’ capacities for shifts in ecologically-relevant morphological traits, like body size and wing length, are an essential aspect of adaptation to changing local conditions (Hoffmann & Sgró 2011). Therefore, when predicting biotic responses to anthropogenic global change, a nuanced understanding of the trajectories of morphological size and shape across species in a community is an important complement to studies of macroecological changes such as phenology and geographic range.

## Methods

### Specimen and data collection

Since 1978, The Field Museum’s collections personnel and volunteers have operated a salvage operation to retrieve birds that collided with buildings in Chicago, IL, USA during their spring or fall migrations (Fig. S1), resulting in approximately 87,000 bird carcasses of more than 200 species brought to the Field Museum from the Chicago area. All measurements included in this study were made by a single person - David E. Willard - who measured the following morphological characteristics on fresh or thawed carcasses prior to preparation as specimens, which should improve the precision of measurements compared to measurements of live birds or dried specimens: 1) tarsus length and bill length using digital calipers; 2) the length of the relaxed wing using a wing rule; and 3) mass using a digital scale. The carcasses were prepared as specimens, and skull ossification (an indication of age), fat levels, sex (from gonadal inspection) and molt were recorded. Skull ossification (Pyle 1997) enabled aging to Hatch Year (HY) or After Hatch Year (AHY). We filtered the dataset (see Supporting Information for details) to 70,716 individuals from 52 species from 1978-2016. These species are from 11 families and 30 genera of mostly passerines (Table S1). Most species in this dataset breed in boreal or temperate forest or edge habitats, but some species are grassland or marsh specialists, and their winter ranges, habitats, migratory distances, life histories and ecologies are diverse (see *Ecology and Natural History*, Supporting Information).

To test for morphological change through time (eqn 1) and the impacts of environmental and climatic variables on morphology (eqn 2), we used two different modeling approaches. We conducted frequentist linear regressions, with the equation-specific independent variables as well as species and year as fixed effects. We also built mixed-effects models, implemented within a Bayesian framework, treating species as a random effect and accounting for phylogenetic relatedness and auto-correlation of variables through time (these models are presented in the Supporting Information, *Bayesian mixed modeling framework*, for details).

### Characterizing change in body size through time

To quantify intra-specific changes in body size from 1978 – 2016, we compared changes in three indices of body size: tarsus, mass and the first axis of a principle component analysis of tarsus, wing, bill and mass.

We modeled the change in tarsus for all specimens that had data on tarsus, year, sex, age (HY or AHY) and species (*n* = 58,475). We used the group-centered logarithms of tarsus for each species as the dependent variable (the logarithm of each tarsus length was taken, and then data within each species was scaled to have a mean of zero and standard deviation of one). For the fixed effects modeling approach, we used a linear model implemented using the ‘lm’ function in R (R Core Team 2018):

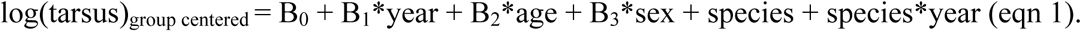

We repeated our analysis of changing body size through time (eqn 1), using log(mass)_group centered_ rather than tarsus as the proxy for body size.

We also conducted a principle components analysis (PCA) of log(tarsus), log(wing length), log(bill length), and log(cube root of mass) for all specimens for which we had data on all measurements (*n* = 48,338) using the ‘princomp’ function in R (R Core Team 2018). Species scores on the first axis of the PCA (PC1) were used as a metric of body size (as is common practice, e.g. (Grant & Grant 2008)). Because all variables were positively loaded onto PC1, and are expected to scale positively with body size, we interpreted PC1 scores as positively related to body size. As with tarsus and mass, we repeated eqn 1 with group centered PC1 scores.

### Change in Wing Length Through Time

Wing length was modeled substituting log(wing length)_group centered_ for tarsus in eqn 1 (*n* = 62,628). In addition to raw wing length, we modeled body size-corrected wing length by regressing log(wing length) onto log(tarsus) for each species (*n* = 58,304) and using the residuals as the dependent variable.

### Environmental Variables

To test hypotheses on the mechanisms underlying changes in body size and wing length, we generated species-specific estimates of climatic and environmental variables (temperature, precipitation, and Normalized Difference Vegetation Index [NDVI], a proxy for resource availability) on the breeding and wintering grounds through time and tested whether they were associated with changes in adult body size. We cropped breeding, wintering and resident ranges for all species (BirdLife International 2015) to exclude unlikely breeding destinations for birds passing through Chicago; we also tested the sensitivity of model results to variations in how ranges were cropped (Supporting Information, Fig. S1). For each species, we then calculated mean temperature, mean precipitation, and maximum mean NDVI through time (1981-2016) in the region representing the likely breeding grounds (June) and on the likely wintering grounds (December) for each species (see Supporting Information).

### Modeling morphology as a function of environmental and climatic variables

To test the impacts of these variables on body size, we modeled tarsus for AHY specimens (HY birds were excluded as they had not experienced winter conditions yet) from 1981 – 2016 (*n* = 29,702). Summer NDVI and summer precipitation were highly correlated (*R* = 0.56), so summer NDVI was not included in the model. The environmental and climate data for the breeding and wintering seasons preceding collection of an individual were used. In order to test whether the relationships between summer variables and body size were similar across both age classes, we modeled the tarsus length of all specimens using eqn 2, but excluding the winter variables, and including age as a predictor.

The analysis of body size as a function of environmental and climatic variables was conducted separately using tarsus or PC1 as the index of body size: body size (i.e. tarsus_*adults, group centered*_ or PC1_*adults, group centered*_) = B_0_ + B_1_* year + B_2_*breeding season precipitation + B_3_*breeding season temperature + B_4_*wintering season precipitation + B_5_*wintering season temperature + B_6_*wintering season NDVI + B_7_*sex + B8*season + species + species*year (eqn 2). Wing length was similarly modeled using eqn 2.

The relative importance of each variable for explaining variance in body size was compared by re-fitting the model across all permutations of model specification and calculating the R^2^ partitioning across those orders (Lindeman *et al*. 1980), implemented using the “calc.relimp” function in the “relaimpo” package in R (Grömping 2006; R Core Team 2018).

To test the sensitivity of our results to uncertainty in AHY age, we compared the results of the tarsus model (eqn 2) to those derived from using the climatic and environmental data from each of the three years preceding collection (Supporting Information).

## RESULTS

### A consistent reduction in body size and increase in wing length in boreal-temperate migratory birds

Despite the ecological and phylogenetic diversity among species, we found consistent reductions in body size across species over the course of the study (Fig. 1, Fig. 2, Fig. 3A, Fig. S2). These reductions in body size were recovered regardless of whether we assessed body size using univariate measurements of either mass or tarsus length, or a multivariate index of size based on the first axis of a principle component analysis of mass, tarsus, wing length, and bill length [PC1]. For simplicity, we present results using tarsus length, as it is the most appropriate proxy of intra-specific body size (Zink, R. M. and Remsen 1986; Rising & Somers 1989; Senar & Pascual 1997), particularly given the extreme variability of mass during migration (Supporting Information). However, all results presented are qualitatively identical whether we measure body size as the univariate tarsus length or the multi-variate PC1 (Supporting Information), and whether we use fixed effects or Bayesian mixed effects models that incorporate phylogenetic relatedness (Supporting Information). Across our dataset, tarsus (hereafter, body size) declined significantly through time (*P* < 0.01) and in nearly all species, and these declines were consistent across age and sex classes (Figs 1 and 2A, Fig. S2).

**Fig. 1.**
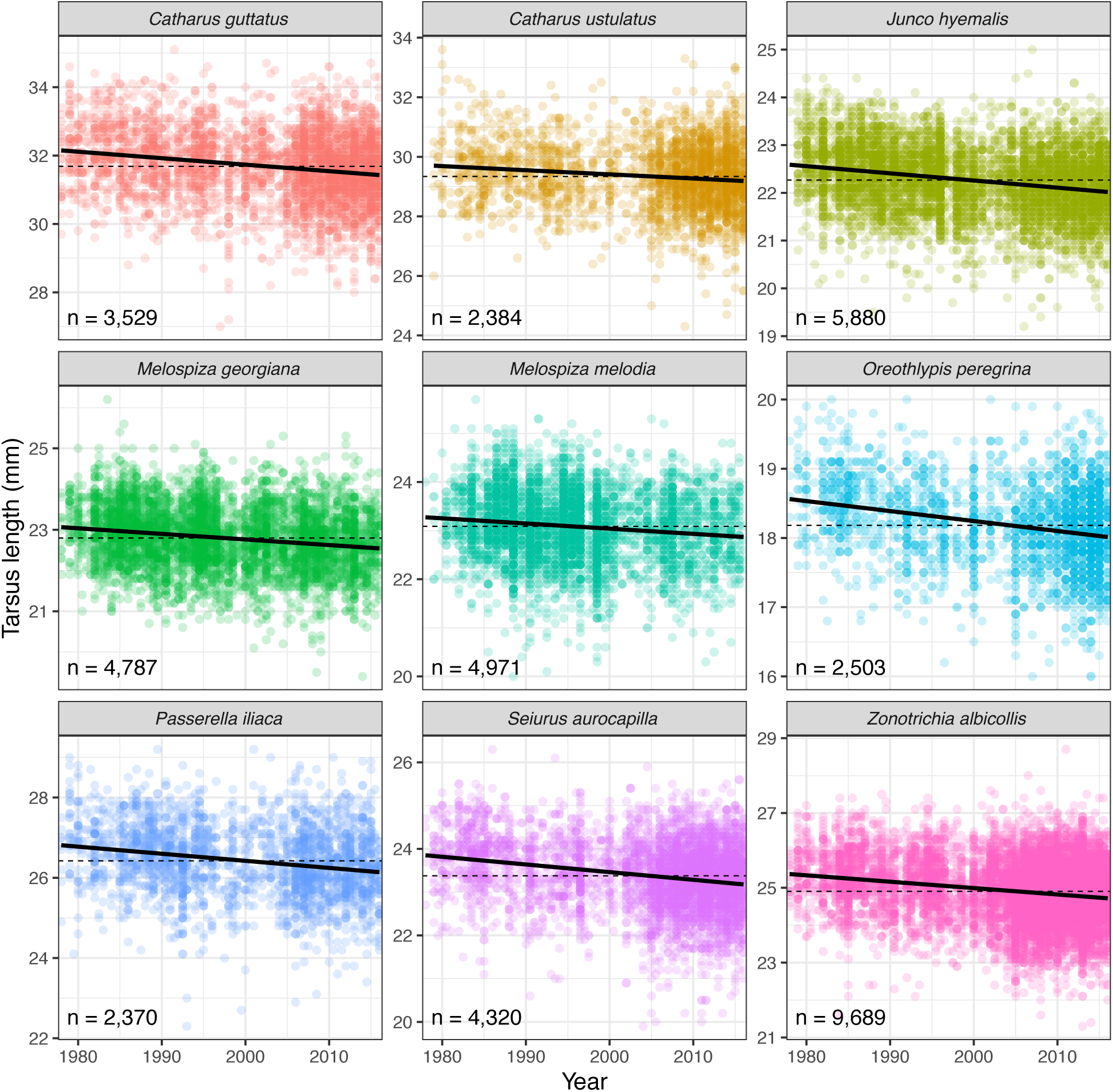
Body size has become smaller through time. Tarsus length declined in nearly all species in the dataset (Fig. 3A) with the 9 most highly sampled species shown here. Dashed lines have a slope of zero and an intercept equal to the mean tarsus length for each species.

**Fig. 2.**
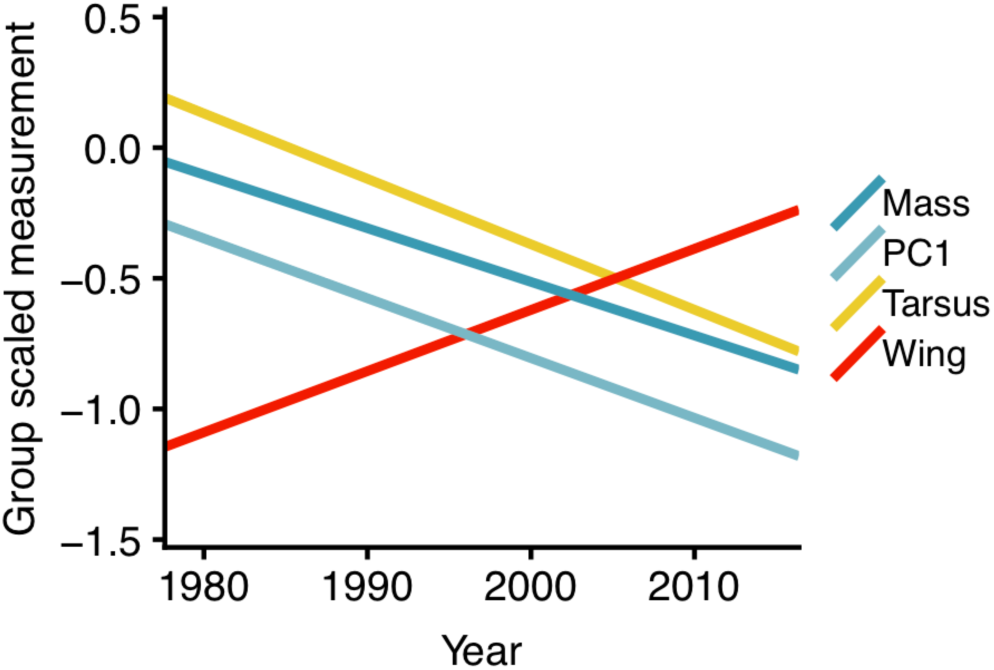
While body size has become smaller, wing length has increased through time. Lines represent all species, with measurements group mean centered by species (70,716 specimens from 52 species). Wing length increased through time (*P* < 0.01), while body size declined (tarsus, mass and the first principal component of a principal components analysis of tarsus, bill, wing and mass all declined through time across species; *P* < 0.01, *P* = 0.056, and *P* < 0.01, respectively).

**Fig. 3.**
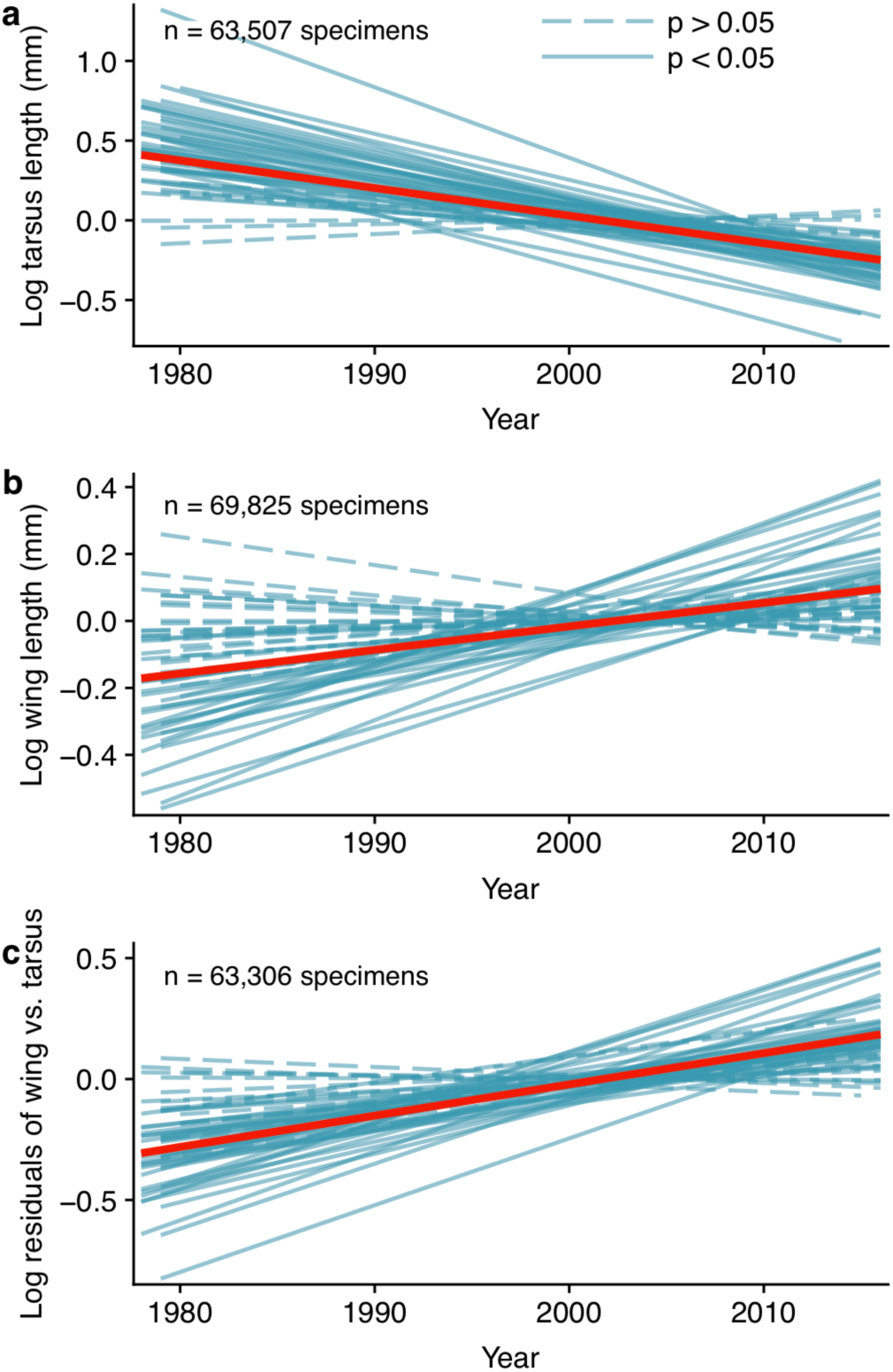
Morphological change and relationships. Measurements are group mean centered by species. Attributes of morphology have changed nearly universally across species (A-C), with individual species trends in blue (slope p-values are shown), and the trend across all species in red (all significantly different from zero). (A) Tarsus has declined in 50/52 species, and all significant changes in tarsus (*P* < 0.05; *n* = 43), represent declines. (B) Wing length has increased through time, and body size-corrected wing length (C) has increased in 47/52 species, and all significant changes (*P* < 0.05; *n* = 35) represent increases.

Body size is positively linearly correlated with wing length (*R* = 0.84 across all species, mean of *R* = 0.28 within species). Nevertheless, as body size declined over time, wing length increased (*P* < 0.01; Fig. 2, Fig. S2). This increase was consistent across all species in our study that showed significant changes in wing length (Fig. 3B). Further, body size-corrected wing length (the residuals of wing length regressed onto body size) similarly increased over the same time period (*P* < 0.001), and this trend was nearly universal (90% of species had increases in relative wing length, and all of the significant changes in relative wing length were positive; Fig. 3C), and was consistent across age and sex classes (Fig. S2). In other words, even those species that have not undergone increases in absolute wing length nevertheless experienced shifts in wing allometry that yielded smaller-bodied, longer-winged birds.

### Increasing summer temperatures drive body size decline

We found that the climatic and environmental variable with the greatest explanatory power for body size—by an order of magnitude—was summer temperature on the breeding grounds, with increased temperatures associated with reduced body size (*P* < 0.001; Table S7). Although various factors beyond temperature, such as food abundance and quality, may contribute to body size reductions (Gardner *et al*. 2011; Sheridan & Bickford 2011; Yom-Tov & Geffen 2011; Teplitsky & Millien 2014), we did not find evidence that proxies for these factors (NDVI and precipitation) have driven the trend in body size.

Although the exact breeding and wintering locations of individuals in the study are not known, as specimens were collected from a passage site, all results are robust to uncertainty in likely breeding locations (Fig. S1). Further, because populations were sampled at a passage site south of the breeding range and north of the wintering range, rather than a single breeding or wintering locality, we are likely collecting individuals from across the latitudinal extent of the species’ ranges, and thus observing broad population-level trends rather than single-site dynamics (Van Buskirk *et al*. 2010).

### Selection during migration drives increases in wing length

The observed increases in wing length were not explained by environmental variables on either the breeding or wintering grounds (Supporting Information). All variables were either not significantly associated with relative wing length (*P* > 0.05), or were significantly associated with wing length but were not changing through time in a way that could produce the observed long-term trend (e.g. a variable may have been significantly positively associated with wing length, but was declining through time; Supporting Information). Additionally, within years, wings were proportionately longer in spring populations than in populations collected during the previous fall migration (*P* < 0.05; Fig. 4B). Notably, in addition to wing length being longer in spring populations, wing length is increasing faster through time in spring birds (Fig. 4B), suggesting selective pressures for increased wing length during migration have been increasing over the course of the study period (see *Discussion*).

**Fig. 4.**
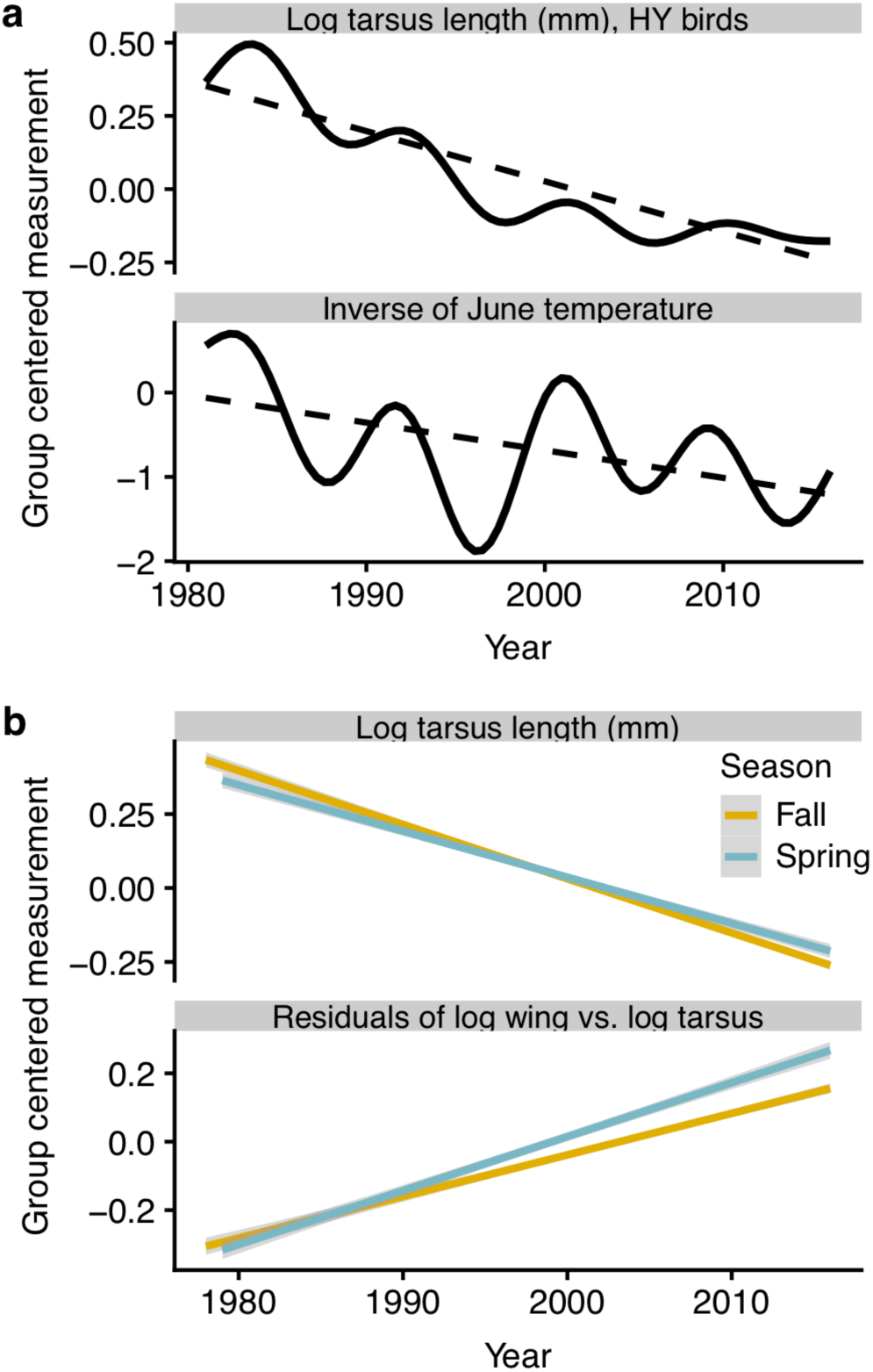
Evidence for temperature-related body size declines and intra-annual selection on wing length. (A) In addition to long term correlated trends in tarsus decline and temperature increase, short term fluctuations in temperature are correlated with short term fluctuations in tarsus length, suggesting a causal relationship in which increasing temperatures are associated with reductions in body size (dashed lines are linear models, solid lines are general additive models). (B) Body size-corrected wing length is longer and is increasing at a more rapid rate in spring birds, reflecting selection for increased wing length during migration.

## DISCUSSION

Despite a diversity of ecologies, habitats, and geographic ranges, we found a near-universal reduction in body size over four decades for the 52 species in our data. The association between temperature and body size recovered by our modeling approach does not reflect merely a long-term correlation between body size and temperature; rather, it also reflects significantly correlated short-term fluctuations after controlling for the long-term trends (Fig. 4A). This result suggests a causal relationship (Methods; (Angrist J. D. and J. S. Pischke 2008)), wherein increasing summer temperatures drive reductions in body size. While other studies have found less consistent reductions in body size in migratory birds (Yom-Tov *et al*. 2006; Salewski *et al*. 2010), this is likely due to the use of mass or wing length as proxies for body size, or smaller sample sizes. Our findings support the hypothesis that body size reduction may be a widespread response to global warming (Gardner *et al*. 2011), occurring broadly across species that tend to be smaller in warmer parts of their range.

Developmental plasticity and selection represent two potential, non-exclusive, mechanisms underlying the observed changes in body size in our data. Experimental studies have shown that increased temperatures during nesting can lead to a reduction in avian adult body size through developmental plasticity (Andrew *et al*. 2017), raising the possibility that the consistent patterns of body size reduction we observe may be a plastic response to increased temperatures during development. Species could also be evolving in response to changing selection pressure on body size. Cold weather metabolic demands are classically invoked to explain Bergmann’s rule (or are considered an integral part of the rule (Watt *et al*. 2010)), with the smaller ratio of surface area to volume that accompanies increased body size considered beneficial in colder climates (Gardner *et al*. 2011; Sheridan & Bickford 2011; Teplitsky & Millien 2014). As such, warming temperatures could conceivably relax selection for larger body size, indirectly leading to size reduction. However, the migratory birds in our study vacate the coldest parts of their ranges during the winter (Winger *et al*. 2018) and also winter in a wide variety of climatic conditions. We found that changes in temperatures on these diverse wintering grounds were not strongly associated with body size changes, suggesting that relaxed cold-season selection pressures on body size are unlikely to explain the observed trends. The observed correlated short-term fluctuations between temperature and body size (Table S7), which were particularly pronounced in hatch year birds (Fig. 4A), suggest a potentially important role for developmental plasticity, particularly given recent experimental evidence for temperature-induced developmental plasticity in body size in passerine birds (Andrew *et al*. 2018). However, it is possible that a combination of developmental plasticity and relaxed selection against smaller body size has yielded the near-universal pattern of body size reduction observed in our data.

More complex ecological dynamics of global change may also contribute to body size reduction, such as food limitation as a result of climate change-driven phenological mismatches (Both *et al*. 2006). Given the observational nature of our data, it is not possible to completely rule out alternative, non-climatic selective pressures (e.g. reduced food availability), particularly if these processes are themselves driven by cyclical fluctuations in temperature. However, because the relationship between temperature and body size is evident after controlling for the long-term trends in the data, an alternative mechanism would need to exhibit both a 40-year correlation with body size as well as correlated short-term fluctuations matching those of body size (Fig. 4A). Further, the near-universality of the morphological changes across the species in our study — which are ecologically diverse and breed and winter in a wide variety of habitats with different phenological dynamics — supports a role for fundamental metabolic or physiological processes influencing the observed trends.

Why has relative wing length increased as body size has declined in nearly all 52 species in our study? In our model results, no climatic or environmental variables on the breeding or wintering grounds explained the long-term increase in wing length (Supporting Information). Together with our finding that spring birds have longer wings than fall birds and that this seasonal difference is widening through time, these results suggest that positive selection for longer relative wings is occurring during migration. These seasonal differences in wing length are likely driven in part by selection on hatch-year birds, which, in many species, tend to have shorter wings [Fig. S2, 68]. Such a pattern of longer wings in spring versus fall could thus alternatively be explained by elevated mortality rates for hatch-year birds that is unrelated to selection on their shorter wing length. However, not only do we find that wing length is longer in spring migrants than fall migrants, but this seasonal difference is increasing through time (Fig. 4B), and wing length is also increasing through time across all age classes (Fig. S2). We interpret the total evidence of these patterns to be indicative of a selective advantage for longer wings during migration that has been increasing over the study period.

Longer and more pointed wings are associated with more efficient flight in birds, particularly for long distance flights such as during migration (Pennycuick 2008), suggesting that some aspect of recent global change is selecting for more efficient flight across this diverse set of migratory birds. Indeed, several global change dynamics have been proposed as mechanisms that should select for increased wing length in migratory birds. These mechanisms include increasing migratory distances associated with poleward range shifts (Förschler & Bairlein 2011), phenological advances requiring faster migrations (Hahn *et al*. 2016; Møller *et al*. 2017), and habitat fragmentation that could require individuals to make longer flights between stopover sites or disperse further to find breeding territories (Desrochers 2010).

Increasing selection for proportionately longer wings during the migratory period could be a result of increasing migratory distance through time. Migratory distance is positively correlated with wing length both within and across species in passerines (Winkler & Leisler 1992; Förschler & Bairlein 2011), suggesting that increases in relative wing length through time could be a response to northward shifts in breeding ranges if wintering ranges have remained static. However, trajectories of warming-induced range shifts have been idiosyncratic across North American bird species (Tingley *et al*. 2009; Mayor *et al*. 2017), while the observed increase in wing length is remarkably consistent across the species in our dataset. Additionally, our data should be robust to changes in geographic distribution, as has been noted in other studies using migratory samples to examine morphological change (Van Buskirk *et al*. 2010). All individuals sampled in our study are from populations that breed north of Chicago and winter south of Chicago, meaning that individuals from across the latitudinal breadth of the breeding grounds (Fig. S1) are likely to have been sampled in Chicago. As such, the majority of our data are likely consistently derived from individuals that breed within the core of their species’ range (Van Buskirk *et al*. 2010), whereas range shifts should lead to selection for longer relative wing lengths at the southern and northern edges of the range. However, identifying the geographic provenance of individuals in our dataset, and how these may have changed through time, will be necessary to directly test the relationship between ranges shifts and morphological change. In addition to investigating how total migratory distances have changed due to latitudinal range shifts, further research should also address the possibility that habitat fragmentation and reduction could select for longer winged individuals (Desrochers 2010) without necessitating a shift of the entire species’ range.

Phenological studies have suggested that migratory birds may be advancing their spring migratory timing in response to climate change (Charmantier & Gienapp 2014). In other studies, birds that migrate earlier and arrive first on the breeding grounds tend to have longer wings than birds that arrive later (Bowlin 2007; Hahn *et al*. 2016). By assuming that passage time through Chicago is correlated with arrival time on the breeding grounds, we tested whether longer-winged birds arrive earlier within years (i.e. does size-corrected wing length predict passage date in a single year; Supporting Information, eqn 3). Our data indicate that longer-winged (*P* < 0.01) and larger (*P* < 0.05) birds do indeed migrate through Chicago earlier in spring than shorter-winged and smaller individuals. However, mean spring passage time through Chicago did not become earlier across years (*P* = 0.31), as would be expected if advancing phenology had selected for increasing wing length through time (Supporting Information). Therefore, we did not find strong evidence that selection for earlier migrations has driven increases in wing length.

Phenological changes, shifting ranges and habitat fragmentation are all plausible and non-exclusive selection pressures that could increase wing length among species; eliminating these competing hypotheses will require a better understanding of the geographic provenance of individuals through time. However, we suggest that the near-universal change in relative wing length across the ecologically and geographically diverse species in our dataset may be evidence of a more fundamental physiological impact of rapid climate change on migratory birds. Specifically, we hypothesize that increased relative wing length confers a selective advantage as body size declines — even for simply maintaining current migratory patterns — due to decreased metabolic efficiency (increased energy required per unit mass; 48) as individuals get smaller. Increased relative wing length improves flight efficiency by reducing wing loading (Rayner 1988), and may additionally reflect an increase in wing pointedness, which further increases flight efficiency (Bowlin & Wikelski 2008; Pennycuick 2008). That is, we propose longer relative wing length may reflect a compensatory adaptation to counter the consequences of shrinking body size for powered flight in migrants. As expected if relative wing length is increasing to compensate for reductions in body size, species in our dataset that have become smaller at faster rates have also experienced faster increases in relative wing length (*P* < 0.05), though this relationship is sensitive to the modeling approach taken (Supporting Information). The complexities of the physics of flight and their relationship with migration (Alerstam & Lindström 1990; Pennycuick 2008; Møller *et al*. 2017), coupled with the dynamic environmental context of migration as the world changes, preclude definitively identifying a mechanistic link between reductions in body size and an increase in wing length to maintain migration. However, understanding if the observed morphological changes in body size and wing length represent a coupled response to global warming — versus decoupled trends driven by alternate forces — is an important avenue of future research, given the consistency with which body size and wing length have changed across this diverse group of species.

While the increase in relative wing length we identified is likely the result of selection during migration and may facilitate the maintenance of migration, it also carries trade-offs for nearly all aspects of avian life history and ecology. Indeed, the tradeoffs associated with variations in wing length are one of the most fundamental components of avian life history, impacting nearly all aspects of ecology and behavior (Norberg 1990). Thus, the extent to which these migratory birds can continue to adapt to rapid global change via shifting wing proportions remains unknown.

## Conclusions

We identify a significant influence of short-term fluctuations in summer temperature on body size that is consistent with the long-term trends shown across species, providing strong evidence that warming temperatures are driving reductions in body size across biota. Body size reduction is likely to have far-reaching ecological consequences (McGill *et al*. 2006). The concomitant increase in wing length may have similarly expansive ecological implications (Norberg 1990), particularly as the divergent trends in body size and wing length combine to drive a change in shape (i.e. increased relative wing length) that may face opposing constraints. Should size and shape be a coupled response to increasing temperatures, tethered by allometric relationships and with broad ecological impacts, understanding how temperature-driven morphological change interacts with shifting phenology geographic range may be essential for predicting biotic responses to climate change.

## Acknowledgements

We thank the staff, curators and volunteers of the Field Museum, and the Chicago Bird Collision Monitors, for their assistance in salvaging birds. For helpful comments, we thank S. Dubay, N. Senner, J. Bates, S. Hackett, B. Marks, J. Voight, M. Jain, and M. Zelditch. We thank D. Megahan for Fig. S1.

## Supporting Information

### Supplementary Methods

#### Data Filtering

For the present study, the following records were removed from the dataset prior to analysis: those with no locality information or those from outside the Chicagoland, IL area (considered here to include Cook, DeKalb, DuPage, Kane, Kendall, Lake, McHenry and Will Counties, although more than 98% of specimens were from Cook County); those with no measurement data; those with no collection date recorded and those unidentified to species. Carcasses were kept in −20°C freezers prior to measurement and preparation as museum specimens. We note that freezing specimens can result in reductions of measurements, particularly mass, due to desiccation. For carcass desiccation to have biased our results, freezing times prior to measurement would needed to have increased steadily over the course of our study, and we have no indication that this has occurred in any consistent way. The vast majority of specimens came from the spring and fall migratory periods; fewer then 1,000 specimens were collected from the summer months (June and July) and were removed because they may have been nestlings or fledglings, and fewer than 300 specimens were collected from the winter months (December, January and February) and were also removed as the focus of this study is on migrants passing through Chicago.

To examine temporal trends in morphology across the broadest set of species, we excluded any species with fewer than 100 total specimens or with fewer than 10 specimens with complete measurement data (i.e., measurements for tarsus, wing and mass) in each period 1980 – 1989,1990 – 1999, 2000 – 2009 and 2000 – 2016. The only exceptions to these criteria were the inclusion of *Certhia americana* and *Sphyrapicus varius*, which were each represented by >2,000 specimens but did not have tarsus measurements from the most recent decade.

Given the size of the dataset, some errors in specimen identification or data entry are inevitable, such that most species contained a handful of obviously erroneous measurements. To remove these, we filtered four measurements (tarsus, wing, bill and mass) to nullify any measurement falling outside an interquartile range of 3 for that measurement for each species (box-and-whisker plots typically identify outliers as those falling outside of a more conservative 1.5 interquartile range; we used a broader range so as to remove errors while attempting to retain true outliers). This filtering procedure nullified only 326 out of 257,985 total measurements.

#### Environmental Data

We calculated temperature on the breeding and wintering grounds using the NASA GISS surface temperature anomaly dataset from 1981 – 2016 (Hansen *et al*. 2010). For each species, to calculate the temperature on the breeding range, the mean June temperature anomaly of each year across the cropped breeding range was used; to calculate the temperature on the wintering range, the mean December temperature anomaly of each year across the cropped wintering range was used (S1 Fig). Temperature data were obtained through the Columbia University IRI data library (https://iridl.ldeo.columbia.edu). Precipitation data were obtained from 1981 – 2016 from the Global Precipitation Climatology Project, provided by NOAA/OAR/ESRL PSD, Boulder, Colorado, USA; https://www.esrl.noaa.gov/psd/ (Adler *et al*. 2003) and used to calculate mean June and mean December precipitation across the cropped breeding and winter ranges, respectively. As a metric for primary productivity, we calculated the maximum mean Normalized Difference Vegetation Index (NDVI), obtained from the NOAA Climate Data Record (Vermote *et al*. 2014) and analyzed using Google Earth Engine (Gorelick *et al*. 2017) from 1981 – 2016. To characterize NDVI on the breeding and wintering ranges, we used the maximum mean NDVI across the breeding range of each species in June and across the wintering range in December.

#### Bayesian Mixed Modeling Framework

In order to test the sensitivity of our analyses to our treatment of the phylogenetic non-independence of our data, we conducted analogous models of morphological change through time (eqn 1) and the influence of climatic and environmental variables on tarsus (eqn 2), using a Bayesian mixed model approach.

For all models examining changes in morphology through time (eqn 1), we conducted an analogous model but within a Bayesian framework in which we treated species identity as a random effect that incorporated a phylogenetic variance covariance matrix. We retrieved 1,000 of the most likely phylogenies for our species from the posterior distribution of a global phylogeny of the birds of the world (https://birdtree.org (Jetz *et al*. 2012)), and calculated a 50% majority rule consensus with branch lengths, following Rubolini et al. (2015) (Rubolini *et al*. 2015). All tips were represented in the phylogeny with genetic data.

Bayesian regression models analogous to the linear model structures we described above (eqn 1) were fit using “brms” (Bürkner 2017) in R (R Core Team 2018). We modeled both the phylogenetic covariance among species and included a parameter to account for species-specific effects not captured in their phylogenetic relatedness. Aside from specifying uninformative prior distributions for the independent variable parameter estimates (normal distribution, mean of 0, standard deviation of 10), brms default prior settings were used. To fit each model, four independent chains were run for 10,000 iterations with the first 1,000 discarded as burn-in; convergence was assessed by examining the posterior distributions of parameter estimates, trace plots and 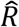 values (with 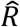 values of 1 considered to reflect convergence).

Similarly, a Bayesian regression model was used to assess the relationship between environmental and climatic variables and body size (eqn 2). In this model, we incorporated phylogenetic relatedness (with the phylogeny constructed as described above) and treated the data as a time series by modeling temporal auto-correlation within brms (using an autoregressive order of 1).

The signs and significance values (whether the significance of a parameter was above or below a threshold of *P* = 0.05) of all parameters were compared to those derived from the linear models (eqns 1 – 2).

### Results

#### Sample Sizes

After applying our species selection criteria, our dataset included 70,716 specimens from 52 species that span 11 families and 30 genera (S1 Table). There was a mean of 1,360 specimens per species, with a range of 101-9,953 (S1 Table). Wing length was measured for 69,825 of the specimens, tarsus was measured for 63,511 specimens, and both wing length and tarsus were measured for 63,306 specimens. Skull ossification was used to specimens collected during the fall to either hatch year (HY) of after hatch year (AHY), and all spring birds were, by definition, characterized as AHY. The dataset contained 67,352 aged birds (32,873 HY; 34,479 AHY).

#### Ecology and Natural History

The only non-passerines were *Porzana carolina* (Rallidae) and *Sphyrapicus varius* (Picidae). The majority of species in the dataset are boreal forest species with breeding ranges either entirely or mainly north of Chicago (e.g., *Zonotrichia albicollis*). However, the dataset also includes some species whose breeding ranges extend further south to encompass Chicago (e.g., *Spizella pusilla*), but whose individuals must have come from north of Chicago. Breeding habitat among the species is diverse, ranging from subarctic taiga (e.g., *Catharus minimus, Spizelloides arborea*) to eastern broadleaf forest (e.g., *Piranga olivacea, Hylocichla mustelina*) to marsh habitats (e.g., *Cistothorus palustris*), edge (e.g., *Passerina cyanea*) or grasslands (e.g. *Ammodramus savannarum*). The wintering ranges and habitats are also diverse, ranging from species in which all individuals winter in South America (e.g., *Setophaga striata, Oporonis agilis*), to those species whose winter ranges include Chicago (e.g., *Junco hyemalis, Spizelloides arborea*) but in which the sampled individuals must have originated south of Chicago. The species are also diverse in diet and foraging strategy; most species are principally insectivorous in the breeding season, but some adopt a more diverse diet in the winter including granivory or frugivory. The species are also diverse in nesting biology, ranging from ground nesters to canopy nesters. Most species build open cup nests, but the dataset also includes some species that nest in cavities or crevices (*Troglodytes aedon* and *Troglodytes hiemalis*) or build covered nests (e.g., *Seiurus aurocapilla*).

#### Body Size Declined Through Time

All indices of body size (tarsus, mass, and PC1) declined through time. Tarsus declined significantly through time (*P* < 0.01), controlling for age, sex, species effects, and species by year interactions (S2 Table). The tarsus model (eqn 1), was significantly better than the null model (*n* = 58,475, *F* = 73.81, *DF* = 105 and 58,369, *P* ≪0.001, adjusted *R*^*2*^ = 0.12). Similarly, mass declined through time, though the relationship is only marginally significant (*P =* 0.056), controlling for age, sex, species effects, and species by year interactions (S3 Table). The mass model (eqn 1) was significantly better than the null model (*n* = 52,390, *F* = 97.95, *DF* = 105 and 52,284, *P* ≪0.001, adjusted *R*^*2*^ *=* 0.16).

The principal component analysis (PCA) of all species with data on wing length, tarsus, bill length, age, sex, and species (*n* = 48,338), had four axes, the first of which (PC1) explained 82% of the variance, with positive loadings on log(wing length) (0.53), log(tarsus) (0.51), log(bill length) (0.44), and log(mass^1/3^) (0.52). The second, third, and fourth axes captured the contrasts between the variables, with inconsistent signs across the loadings for the variables. PC1 declined through time (indicating body size has declined through time), and this decline was significant (*P* < 0.01) after controlling for age, sex, species effects and species by year interactions (S4 Table). This decline is particularly notable given the expectation that increasing temperatures should drive increasing relative bill and, to a lesser degree, tarsus length (Allen’s rule (Symonds & Tattersall 2010)). The model was significantly better than the null model (*n* = 48,338, *F* = 284.8, *DF* = 105 and 48,232, *P* ≪0.001, adjusted *R*^*2*^ = 0.38).

Given the significant decline in tarsus and PC1, and the near-significant decline in mass, our interpretation is that overall body size has declined through time. Tarsus is a better indicator of intraspecific body size in passerines than wing length (Rising & Somers 1989; Senar & Pascual 1997). Mass is expected to have higher variance given rapid fat gains and losses of migratory birds in migration (Alerstam & Lindström 1990; Morris *et al*. 1996), so it is not surprising that the mass trend was consistent with the tarsus and PC1 trends, but less statistically significant. Although estimates of body size derived from multivariate principal components analyses are often desirable, we focus on tarsus as an indicator of intraspecific changes in body size, as it is not as vulnerable to fluctuations in mass (either induced by actual variations in mass that occur during migration or as a result of dehydration of specimens prior to measurement) that may impact changes in PC1.

#### Wing Length Increased Through Time

Raw wing length increased significantly through time (*P* < 0.01), controlling for age, sex, species effects, and species by year interactions (S5 Table). The model was significantly better than the null model (*n* = 62,628, *F* = 496, *DF* = 105 and 62,522, *P* ≪0.001, adjusted *R*^*2*^ *=* 0.45)

Similarly, relative wing length increased significantly through time (*P* < 0.001), controlling for age, sex, year, species effects, and species by year interactions (S6 Table). The model was significantly better than the null model (*n =* 58,304, *F* = 379.8, *DF* = 105 and 58,198, *P* ≪0.001, adjusted *R*^*2*^ = 0.41).

In addition to the long-term trends in relative wing length, we modeled the effect of season on relative wing length, controlling for time, season, sex, species effects, and species by year interactions. The model was significantly better than the null model (*n* = 58,304, *F* = 366.6, *DF* = 105 and 58,198, *P* ≪0.001, adjusted *R*^*2*^ *= 0*.*4)*. In this model, spring had a positive and significant (*P* < 0.05) relationship with relative wing length.

#### Climatic and Environmental Predictors of Tarsus

We modeled body size as a function of climatic and environmental predictors for AHY birds from 1981-2016 (eqn 2), using both tarsus and PC1 as the index of body size. Precipitation on the breeding grounds and NDVI on the breeding grounds were highly correlated (*R* = 0.56), so NDVI on the breeding grounds was not included in the model. Of the variance explained by the model (amounts of variance explained are for tarsus, followed by PC1), the variables that contributed the most were sex (68%, 70%), year (22%, 24%), temperature on the breeding grounds (3%, 2%), species by year interactions (3%, 1%), and species effects (2%, 2%; this effect is small because the data were group-mean centered by species). Both tarsus and PC1 were significantly larger in males (*P* < 0.001), declined through time (*P* < 0.05), and was significantly negatively associated with temperature on the breeding grounds (*P* < 0.001). The remaining climatic and environmental variables each explained less than 1% of the variance explained by the models. The tarsus model was significantly better than the null model (*n* = 29,702, *F* = 37.41, *DF* = 110 and 29,591, *P* ≪0.001, adjusted *R*^*2*^ = 0.12), as was the PC1 model (*n* = 24,012, *F* = 137.5, *DF* = 110 and 23,901, *P* ≪0.001, adjusted *R*^*2*^ = 0.38)

In addition to modeling the impacts of both summer and winter variables on size, we modeled tarsus for all specimens, including both HY and AHY birds, using eqn 2, without any of the winter variables (as the HY birds had not yet lived through a winter season), and with the addition of age as a covariate. The results were qualitatively similar to the model that only included adult birds, with the most variance explained by sex (68%), year (24%), species effects (3%), temperature on the breeding grounds (2%), and species by year interactions (2%). All other variables, including age, explained less than 1% of the variance explained by the model. The model was significantly better than the null model (*n* = 57,718, *F* = 69.64, *DF* = 108 and 57,609, *P ≪*0.001, adjusted *R*^*2*^ = 0.11).

#### Climatic and Environmental Predictors of Wing Length

The model of wing length of AHY birds as a function of climatic and environmental variables was (eqn 2) was significantly better than the null model (*n* = 31,987, *F* = 253.6, *DF* = 110 and 31,876, *P* ≪0.001, adjusted *R*^*2*^ = 0.46). Uncorrected wing length increased significantly through time (*P* < 0.01), and was significantly positively associated with winter NDVI (*P* < 0.01) and winter precipitation (*P* < 0.001). Despite the significant association, winter precipitation cannot explain the long-term increase in wing length, as winter precipitation has significantly declined through time (*P* < 0.001) but was positively associated with wing length. Winter NDVI is positively associated with wing length, and has significantly increased through time, making it a potential driver of the long-term trend in wing length. However, winter NDVI explained less than 1% of the variance explained by the model, suggesting it is not contributing to the long-term change in wing length. More generally, with the exception of year, which explained 2% of the variance explained by the model, no environmental or climatic variables explained more than 1% of the variance explained by the model.

The most variance in body size-corrected wing length was explained by sex (88%), year (6%), season (4%), and species by time interactions (1%). All environmental and climatic variables, with the exception of winter temperature, were significantly associated with relative wing length (*P* < 0.05), but they all explained less than 1% of the variance explained by the model.

#### Results Using a Bayesian Mixed Modeling Framework and Phylogenetic Correction

All parameter estimates converged, with 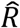 values of 1. The relationship between year tarsus, mass, PC1, wing length, and body size-corrected wing length through time were qualitatively similar (in sign) in the Bayesian models and the linear fixed effect model results (eqn 1). The only differences in statistical significance across the models was a significant relationship between mass and year in the Bayesian model, while that relationship was only marginally significant (*P =* 0.056) in the fixed effects linear models.

The relationships between environmental and climatic factors and AHY tarsus length were also qualitatively similar in both the frequentist fixed effects model (eqn 2) and the analogous Bayesian mixed effects model. All parameter estimates had converged, with ! values of 1. All parameter signs were the same across modeling frameworks. All relationships and significance values were similar in sign and significance when the relationships between environmental and climatic variables (eqn 2) and tarsus length was modeled for all birds (including HY birds) and only summer variables, except that the association with precipitation on the breeding ground changes from marginally significant to significant.

#### Arrival Time

In order to test the influence of body size and wing length on arrival time within years, and shifts in arrival time across years, we modeled arrival time for individuals collected during their spring migration from 1979 – 2016. We filtered out any species that did not have arrival data from at least ten years, after removing any years in which specimens from that species were collected on fewer than five days. This left 26 species with data from at least ten years in which specimens of that species were collected on at least five days (*n* = 19,652).

In order to test for the impact of body size on arrival time within years, we used the within-year collection date: collection date = B_0_ + B_1_*tarsus_group centered_ + species + species*tarsus (eqn 3). Similarly, to measure the effects of relative wing length on with-year arrival time, we fit eqn 3 using body size-corrected wing length, rather than tarsus.

In order to test for shifts in the arrival time across years, we modeled within-year collection date (again, scaled to have a mean of zero and a variance of one): collection date = B_0_ + B_1_*year + species + species*year (eqn 4).

Within-year collection date was significantly negatively associated with tarsus (i.e. larger birds arrived earlier; *P* < 0.01), and the model (eqn 3) was significantly better than a null model (*F* = 1,688, *DF* = 52 and 19,599, *P ≪*0.001, *R*^*2*^ = 0.82). Within-year collection date was similarly significantly negatively associated with group centered relative wing length (*P* < 0.01), and the model (eqn 3) was significantly better than a null model (*F* = 1,823, *DF* = 52 and 19,599, *P* <<0.001, *R*^*2*^ *=* 0.83). Across years, collection date has not changed significantly (eqn 4; *P* = 0.31).

#### Rates of Change in Tarsus Predict Rates of Change in Wing Length

For each species, we modeled group-centered tarsus and body size-corrected wing length through time for each species. We retained the slope of the model for each species as well as the variance of the slope parameter estimate. In order to test the hypothesis that increases in size-corrected wing length are associated with reductions in body size, we modeled the rate of change in size-corrected wing length as a function of the rate of change in tarsus length. (*n* =52). The uncertainty in the slope estimates was treated as measurement error, and phylogenetic correlation was accounted for using the “GLSME” function in the GLSME package in R (Hansen & Bartoszek 2012); our results were sensitive to our treatment of bias.

Because of the variable slopes in our data, and the different levels of error across slopes and variables, we corrected for bias despite a low reliability ratio (Hansen & Bartoszek 2012). Significance of the parameter estimate was assessed based on whether the distance of the parameter estimate from zero was more than twice the estimated standard error of parameter (Gelman & Hill 2009). Slope in wing length through time was negatively related to slope in tarsus through time (i.e. those species with greater rates of loss in tarsus experienced greater rates of increase in relative wing length; *n =*52). The bias-corrected GLS parameter estimate (Hansen & Bartoszek 2012) was −1*10^−4^, which was more than twice the standard error in the bias parameter estimate (2*10^−19^), suggesting the parameter value is significantly different from zero (Gelman & Hill 2009). Importantly, this result was sensitive to our decision to correct for bias within the error structure of the measurement error; the parameter estimate, when not correcting for bias, was not significant.

#### Sensitivity of Results to Time Lag

The relationships between temperature on the breeding grounds and tarsus length was robust to our treatment of year, despite not knowing the exact age of AHY birds. Tarsus was significantly negatively related to temperature on the breeding grounds across all models. With the exception of the three-year lag, temperature on the breeding grounds was consistently responsible for explaining more of the variance explained by the model than any other environmental or climatic predictor, and in two of the three models, it explained an order of magnitude more variance than the next most important predictor. In the one treatment of time in which summer breeding temperature was not the most significant predictor (when climate and environmental data from three years earlier was used), the most important variable was precipitation on the wintering grounds. Winter precipitation could not explain the long-term trend in tarsus, as winter precipitation was positively related to tarsus, but increased through time while tarsus declined.

### Supplemental Figures

**Supplementary Figure 1.**
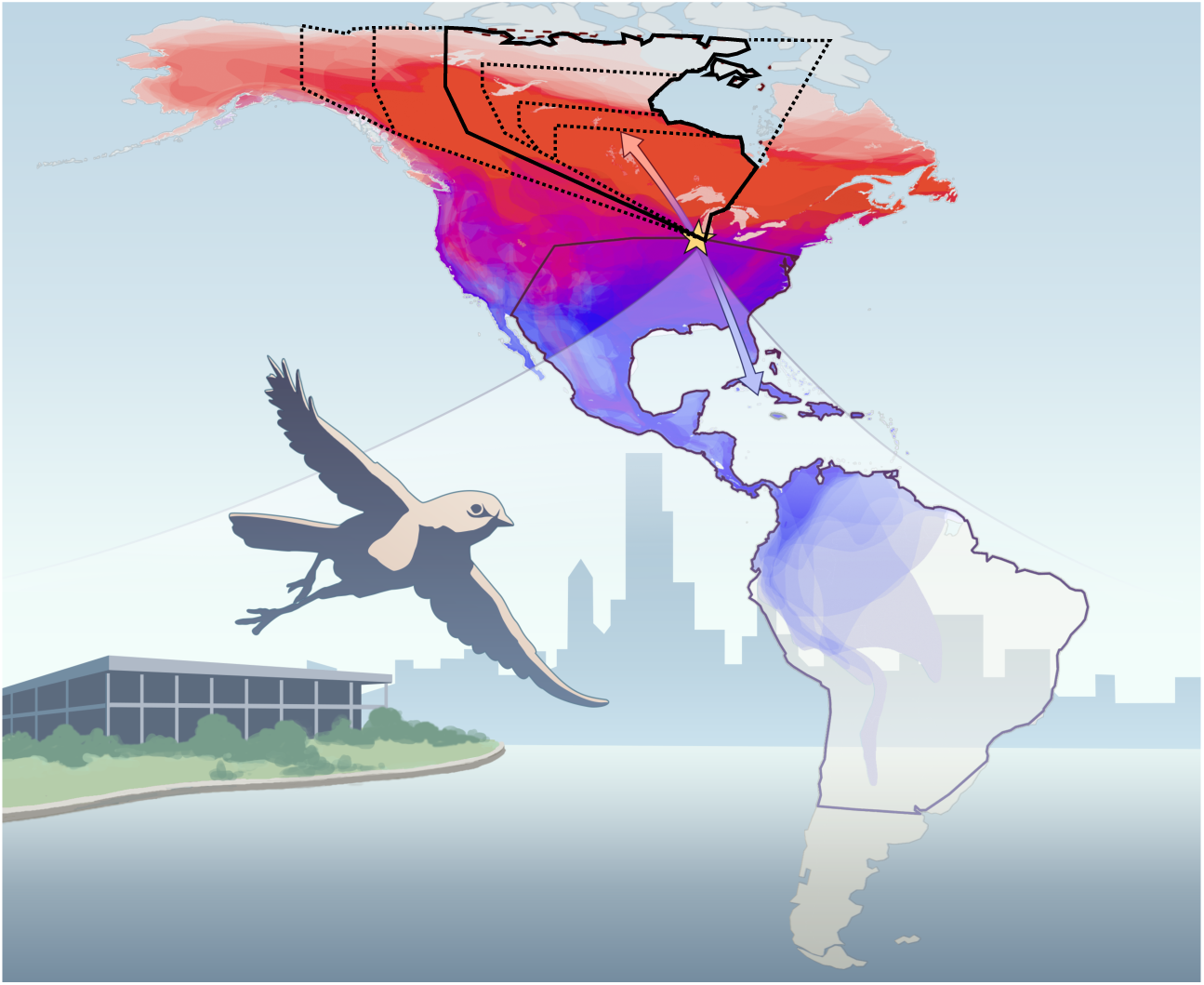
Data collection and sensitivity to Subsetting of breeding ranges. All individuals included in the study were collected after they collided with buildings in Chicago, IL during fall or spring migration. The species’ breeding ranges span North America (individual species’ breeding ranges are outlined in red) and winter ranges extend from the southern United States through the Neotropics (individual species’ wintering ranges are outlined in blue). Likely destinations (solid and dashed lines) were determined based on known migratory paths, and environmental and climatic variables were calculated for the intersection of each species’ range and their likely destinations; modeling results were robust to how these regions were defined. We modeled the relationship between body size and environmental variables (eqn 2) using different subsets of the breeding ranges of each species to calculate the environmental variables. The model results reported in the text are based on the region outlined with the solid line. We found similar results – temperature had a significant negative relationship with body size, and explained the most variance of any variable – using all areas outlined in dashed lines.

**Supplementary Figure 2.**
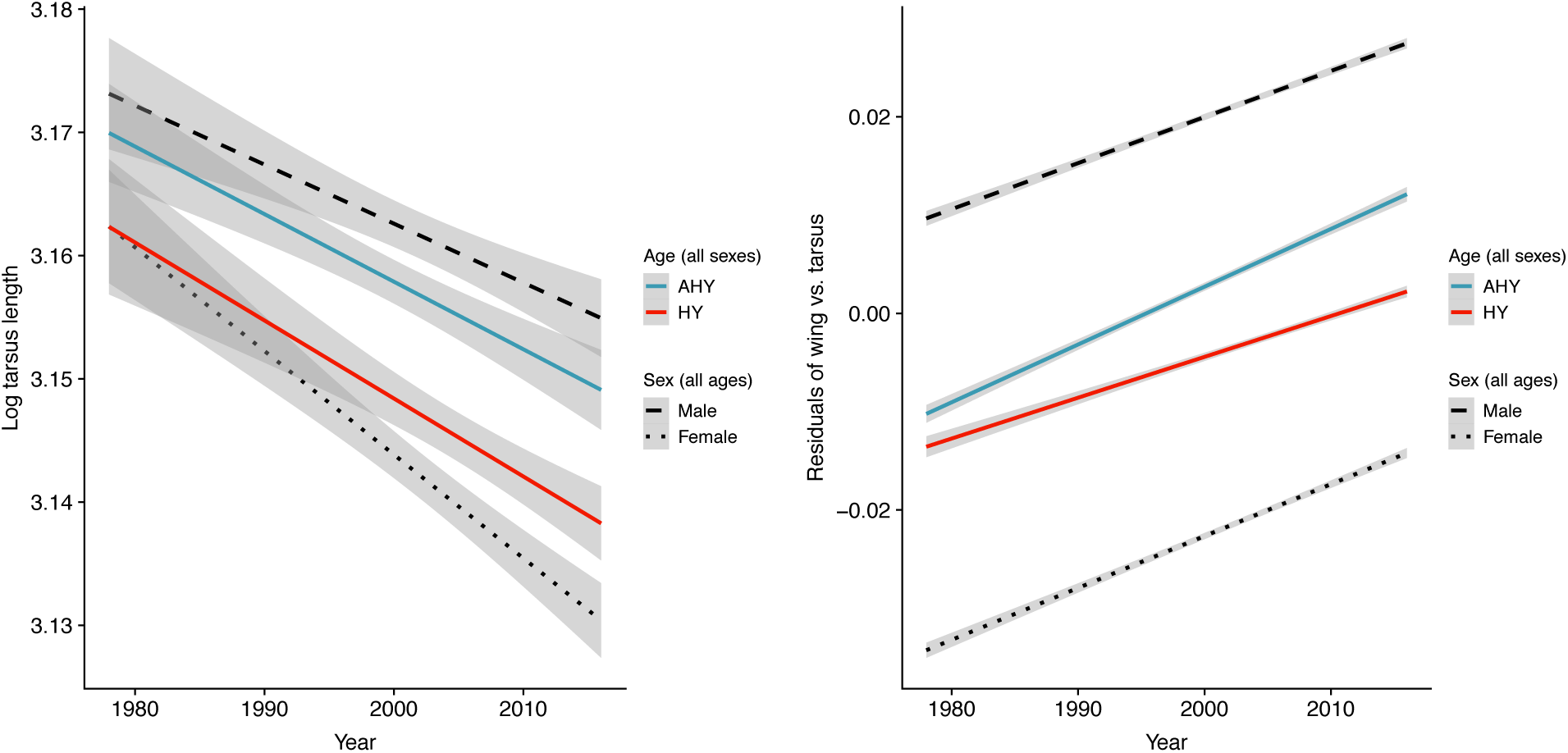
Body Size has Declined and Relative Wing Length has Increased Across Age and Sex Classes. While relative wing length has increased in both age classes (right), this increase is more pronounced in adult (AHY) birds. This is consistent with selection for increased wing length during migration as a mechanism for long-term increases in wing length, rather than simple intra-annual shifts in demography.

### Supplemental Tables

**Supplementary Table 1.**
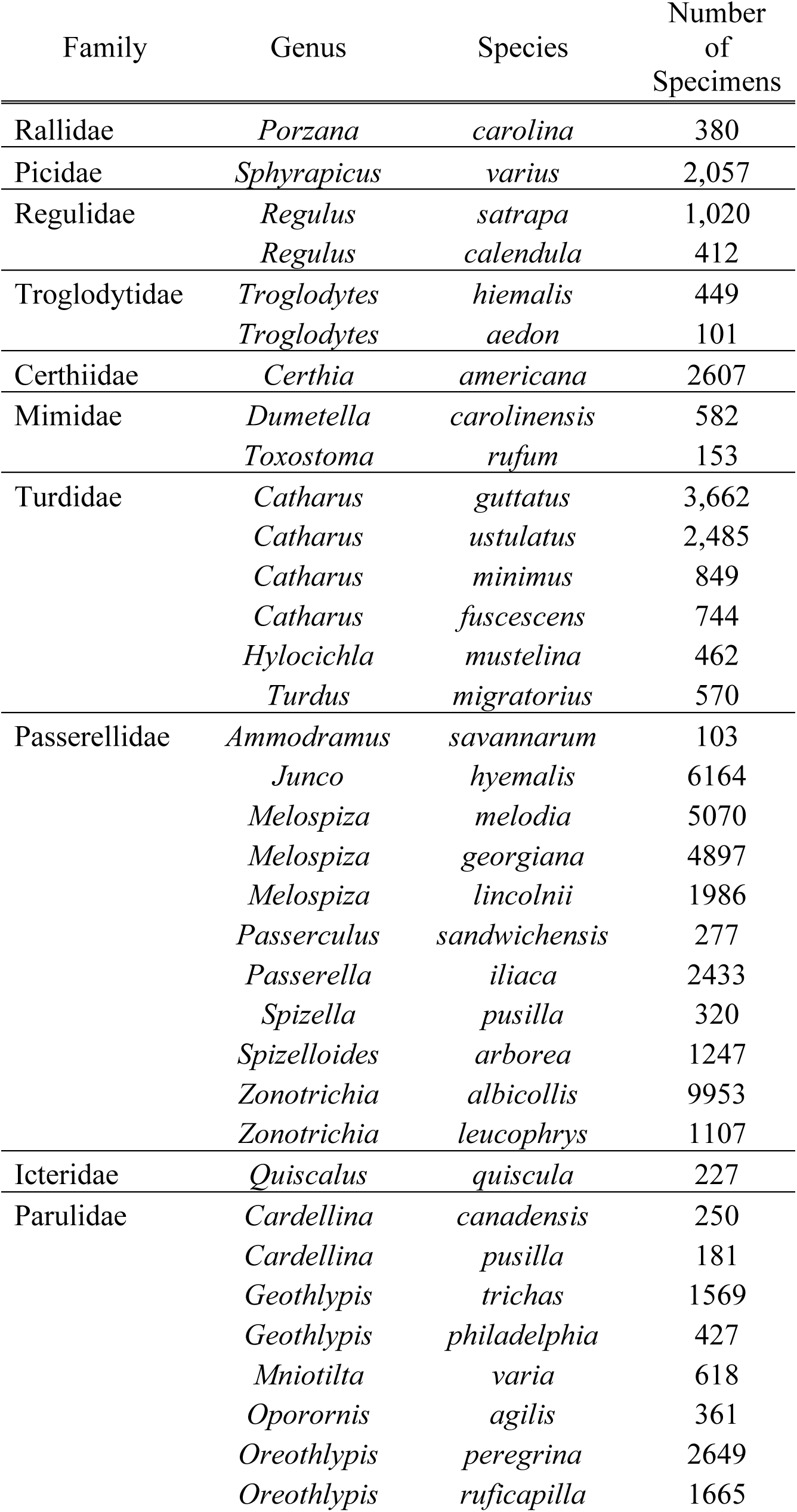
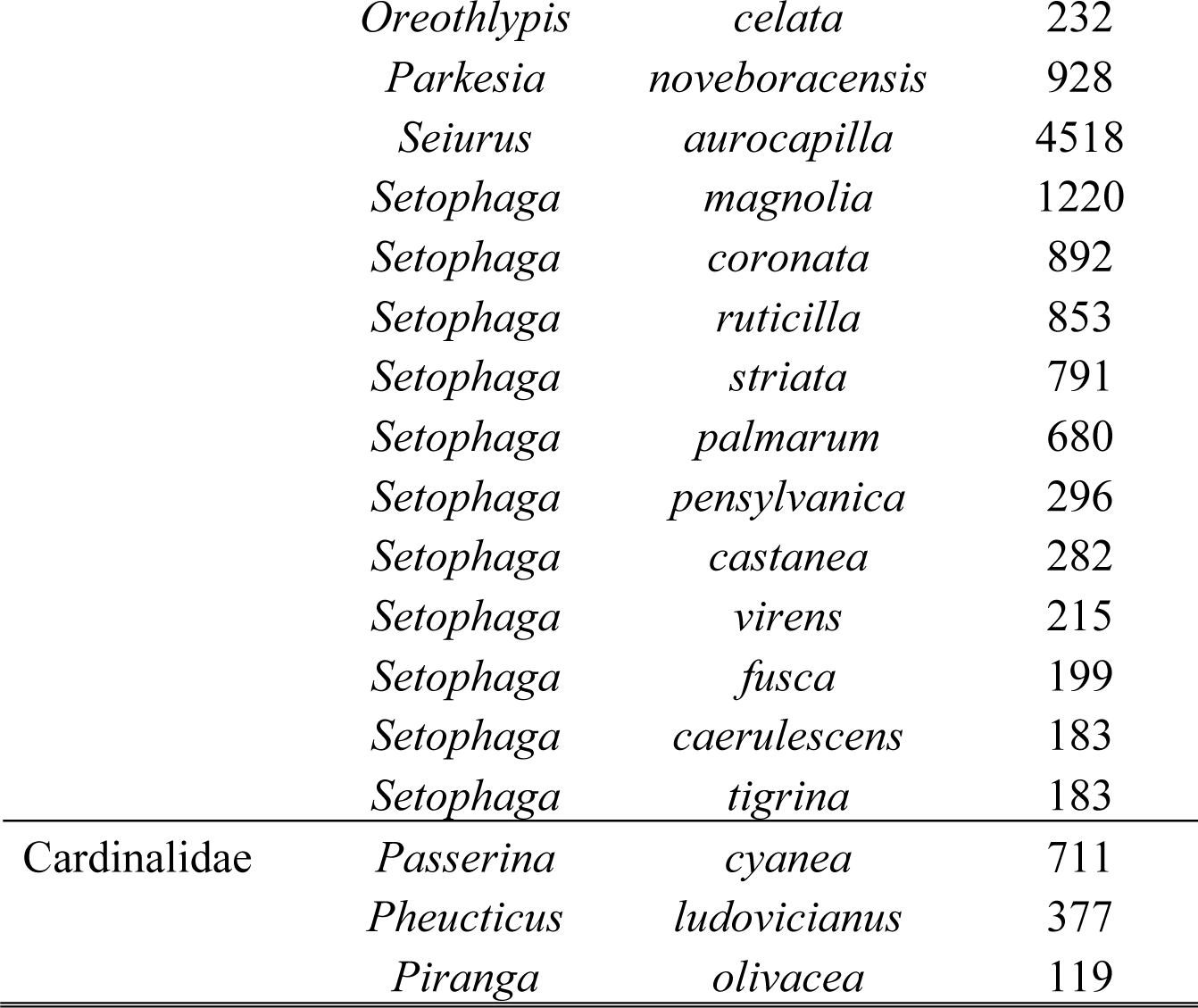
*Taxonomic Sampling in the Dataset*. After filtering the data (*Materials and Methods*), the dataset included 70,716 specimens from 52 species spanning 11 families and 30 genera.

**Supplementary Table 2.** *Tarsus Length has Decreased through Time*. All species and species by year interaction terms are relative to the reference taxon, *Ammodramus savannorum*.

| Variable | Parameter Estimate |
| --- | --- |
| Intercept | 53.045*** |
| Year | -0.027*** |
| Age | -0.036*** |
| Sex (male) | 0.557*** |
| Species effects |  |
| *Cardellina canadensis* | -4.4 |
| *Cardellina pusilla* | 15.8 |
| *Catharus fuscescens* | -26.6 |
| *Catharus guttatus* | -19.2 |
| *Catharus minimus* | -22.0 |
| *Catharus ustulatus* | -27.6 |
| *Certhia americana* | 15.5 |
| *Dumetella carolinensis* | -25.3 |
| *Geothlypis philadelphia* | -5.4 |
| *Geothlypis trichas* | -5.1 |
| *Hylocichla mustelina* | -31.0 |
| *Junco hyemalis* | -13.8 |
| *Melospiza georgiana* | -18.5 |
| *Melospiza lincolnii* | -21.3 |
| *Melospiza melodia* | -23.9 |
| *Mniotilta varia* | -2.6 |
| *Oporornis agilis* | -28.1 |
| *Oreothlypis celata* | -12.3 |
| *Oreothlypis peregrina* | -10.1 |
| *Oreothlypis ruficapilla* | -5.3 |
| *Parkesia noveboracensis* | -1.0 |
| *Passerculus sandwichensis* | -36.0 |
| *Passerella iliaca* | -18.2 |
| *Passerina cyanea* | -9.9 |
| *Pheucticus ludovicianus* | -40.0* |
| *Piranga olivacea* | -26.4 |
| *Porzana carolina* | -56.5** |
| *Quiscalus quiscula* | -59.4** |
| *Regulus calendula* | -40.4* |
| *Regulus satrapa* | -30.6 |
| *Seiurus aurocapilla* | -11.8 |
| *Setophaga caerulescens* | -1.6 |
| *Setophaga castanea* | -6.2 |
| *Setophaga coronata* | -13.5 |
| *Setophaga fusca* | 39.5 |
| *Setophaga magnolia* | -4.0 |
| *Setophaga palmarum* | -16.5 |
| *Setophaga pensylvanica* | -18.6 |
| *Setophaga ruticilla* | 0.3 |
| *Setophaga striata* | -9.1 |
| *Setophaga tigrina* | -5.0 |
| *Setophaga virens* | -2.3 |
| *Sphyrapicus varius* | -47.8 |
| *Spizella pusilla* | -8.2 |
| *Spizelloides arborea* | -36.0* |
| *Toxostoma rufum* | -70.1*** |
| *Troglodytes aedon* | -20.0 |
| *Troglodytes hiemalis* | -27.9 |
| *Turdus migratorius* | -27.8 |
| *Zonotrichia albicollis* | -15.1 |
| *Zonotrichia leucophrys* | -23.7 |

|  |  |
| --- | --- |
| Year:*Cardellina canadensis* | 0.002 |
| Year:*Cardellina pusilla* | -0.008 |
| Year:*Catharus fuscescens* | 0.013 |
| Year:*Catharus guttatus* | 0.01 |
| Year:*Catharus minimus* | 0.011 |
| Year:*Catharus ustulatus* | 0.014 |
| Year:*Certhia americana* | -0.008 |
| Year:*Dumetella carolinensis* | 0.013 |
| Year:*Geothlypis philadelphia* | 0.003 |
| Year:*Geothlypis trichas* | 0.003 |
| Year:*Hylocichla mustelina* | 0.016 |
| Year:*Junco hyemalis* | 0.007 |
| Year:*Melospiza georgiana* | 0.009 |
| Year:*Melospiza lincolnii* | 0.011 |
| Year:*Melospiza melodia* | 0.012 |
| Year:*Mniotilta varia* | 0.001 |
| Year:*Oporornis agilis* | 0.014 |
| Year:*Oreothlypis celata* | 0.006 |
| Year:*Oreothlypis peregrina* | 0.005 |
| Year:*Oreothlypis ruficapilla* | 0.003 |
| Year:*Parkesia noveboracensis* | 0.001 |
| Year:*Passerculus sandwichensis* | 0.018 |
| Year:*Passerella iliaca* | 0.009 |
| Year:*Passerina cyanea* | 0.005 |
| Year:*Pheucticus ludovicianus* | 0.020* |
| Year:*Piranga olivacea* | 0.013 |
| Year:*Porzana carolina* | 0.028** |
| Year:*Quiscalus quiscula* | 0.030** |
| Year:*Regulus calendula* | 0.020* |
| Year:*Regulus satrapa* | 0.015 |
| Year:*Seiurus aurocapilla* | 0.006 |
| Year:*Setophaga caerulescens* | 0.001 |
| Year:*Setophaga castanea* | 0.003 |
| Year:*Setophaga coronata* | 0.007 |
| Year:*Setophaga fusca* | -0.019 |
| Year:*Setophaga magnolia* | 0.002 |
| Year:*Setophaga palmarum* | 0.008 |
| Year:*Setophaga pensylvanica* | 0.009 |
| Year:*Setophaga ruticilla* | 0.00003 |
| Year:*Setophaga striata* | 0.005 |
| Year:*Setophaga tigrina* | 0.003 |
| Year:*Setophaga virens* | 0.001 |
| Year:*Sphyrapicus varius* | 0.024 |
| Year:*Spizella pusilla* | 0.004 |
| Year:*Spizelloides arborea* | 0.018* |
| Year:*Toxostoma rufum* | 0.035*** |
| Year:*Troglodytes aedon* | 0.01 |
| Year:*Troglodytes hiemalis* | 0.014 |
| Year:*Turdus migratorius* | 0.014 |
| Year:*Zonotrichia albicollis* | 0.008 |
| Year:*Zonotrichia leucophrys* | 0.012 |
| Observations | 58,475 |
| R<sup>2</sup> | 0.117 |
| Adjusted R<sup>2</sup> | 0.116 |
| Residual Std. Error | 0.940 (df = 58,369) |
| F Statistic | 73.8*** (df = 105; 58,369) |
| *p<0.1, **p<0.05, ***p<0.01 |  |

**Supplementary Table 3.** *Mass has Declined through Time*. All species and species by year interaction terms are relative to the reference taxon *Ammodramus savannorum*.

| Variable | Parameter Estimate |
| --- | --- |
| Intercept | 39.940* |
| Year | -0.020* |
| Age | -0.073*** |
| Sex (male) | 0.782*** |
| Species Effects |  |
| *Cardellina canadensis* | -40.393* |
| *Cardellina pusilla* | -77.162*** |
| *Catharus fuscescens* | -22.669 |
| *Catharus guttatus* | -19.68 |
| *Catharus minimus* | -0.694 |
| *Catharus ustulatus* | -20.413 |
| *Certhia americana* | 11.069 |
| *Dumetella carolinensis* | -55.384** |
| *Geothlypis philadelphia* | -59.840*** |
| *Geothlypis trichas* | -42.655** |
| *Hylocichla mustelina* | -31.793 |
| *Junco hyemalis* | -21.893 |
| *Melospiza georgiana* | -34.813 |
| *Melospiza lincolnii* | -30.026 |
| *Melospiza melodia* | -25.975 |
| *Mniotilta varia* | -39.182* |
| *Oporornis agilis* | -6.563 |
| *Oreothlypis celata* | -24.88 |
| *Oreothlypis peregrina* | -14.333 |
| *Oreothlypis ruficapilla* | -28.319 |
| *Parkesia noveboracensis* | -36.364* |
| *Passerculus sandwichensis* | -41.306 |
| *Passerella iliaca* | -29.888 |
| *Passerina cyanea* | -27.7 |
| *Pheucticus ludovicianus* | -37.289 |
| *Piranga olivacea* | -26.774 |
| *Porzana carolina* | -24.687 |
| *Quiscalus quiscula* | -22.878 |
| *Regulus calendula* | -24.864 |
| *Regulus satrapa* | -4.766 |
| *Seiurus aurocapilla* | -36.539* |
| *Setophaga caerulescens* | -45.315* |
| *Setophaga castanea* | 8.706 |
| *Setophaga coronata* | -0.49 |
| *Setophaga fusca* | -42.113 |
| *Setophaga magnolia* | -33.13 |
| *Setophaga palmarum* | -13.173 |
| *Setophaga pensylvanica* | -38.026 |
| *Setophaga ruticilla* | -40.986* |
| *Setophaga striata* | -3.937 |
| *Setophaga tigrina* | -4.867 |
| *Setophaga virens* | -47.004* |
| *Sphyrapicus varius* | -18.083 |
| *Spizella pusilla* | -20.458 |
| *Spizelloides arborea* | -34.536 |
| *Toxostoma rufum* | 3.564 |
| *Troglodytes aedon* | -20.06 |
| *Troglodytes hiemalis* | 6.796 |
| *Turdus migratorius* | -27.86 |
| *Zonotrichia albicollis* | -30.804 |
| *Zonotrichia leucophrys* | -38.853* |

|  |  |
| --- | --- |
| Year:*Cardellina canadensis* | 0.020* |
| Year:*Cardellina pusilla* | 0.039*** |
| Year:*Catharus fuscescens* | 0.011 |
| Year:*Catharus guttatus* | 0.01 |
| Year:*Catharus minimus* | 0.0004 |
| Year:*Catharus ustulatus* | 0.01 |
| Year:*Certhia americana* | -0.005 |
| Year:*Dumetella carolinensis* | 0.028** |
| Year:*Geothlypis philadelphia* | 0.030*** |
| Year:*Geothlypis trichas* | 0.021** |
| Year:*Hylocichla mustelina* | 0.016 |
| Year:*Junco hyemalis* | 0.011 |
| Year:*Melospiza georgiana* | 0.017 |
| Year:*Melospiza lincolnii* | 0.015 |
| Year:*Melospiza melodia* | 0.013 |
| Year:*Mniotilta varia* | 0.020* |
| Year:*Oporornis agilis* | 0.003 |
| Year:*Oreothlypis celata* | 0.013 |
| Year:*Oreothlypis peregrina* | 0.007 |
| Year:*Oreothlypis ruficapilla* | 0.014 |
| Year:*Parkesia noveboracensis* | 0.018* |
| Year:*Passerculus sandwichensis* | 0.021* |
| Year:*Passerella iliaca* | 0.015 |
| Year:*Passerina cyanea* | 0.014 |
| Year:*Pheucticus ludovicianus* | 0.019 |
| Year:*Piranga olivacea* | 0.013 |
| Year:*Porzana carolina* | 0.012 |
| Year:*Quiscalus quiscula* | 0.011 |
| Year:*Regulus calendula* | 0.012 |
| Year:*Regulus satrapa* | 0.003 |
| Year:*Seiurus aurocapilla* | 0.018* |
| Year:*Setophaga caerulescens* | 0.023* |
| Year:*Setophaga castanea* | -0.004 |
| Year:*Setophaga coronata* | 0.0003 |
| Year:*Setophaga fusca* | 0.021 |
| Year:*Setophaga magnolia* | 0.017 |
| Year:*Setophaga palmarum* | 0.007 |
| Year:*Setophaga pensylvanica* | 0.019 |
| Year:*Setophaga ruticilla* | 0.021* |
| Year:*Setophaga striata* | 0.002 |
| Year:*Setophaga tigrina* | 0.003 |
| Year:*Setophaga virens* | 0.024* |
| Year:*Sphyrapicus varius* | 0.009 |
| Year:*Spizella pusilla* | 0.01 |
| Year:*Spizelloides arborea* | 0.017 |
| Year:*Toxostoma rufum* | -0.002 |
| Year:*Troglodytes aedon* | 0.01 |
| Year:*Troglodytes hiemalis* | -0.003 |
| Year:*Turdus migratorius* | 0.014 |
| Year:*Zonotrichia albicollis* | 0.016 |
| Year:*Zonotrichia leucophrys* | 0.020* |
| Observations | 52,390 |
| R<sup>2</sup> | 0.164 |
| Adjusted R<sup>2</sup> | 0.163 |
| Residual Std. Error | 0.915 (df = 52,284) |
| F Statistic | 97.954*** (df = 105; 52,284) |
| *p<0.1, **p<0.05, ***p<0.01 |  |

**Supplementary Table 4.**
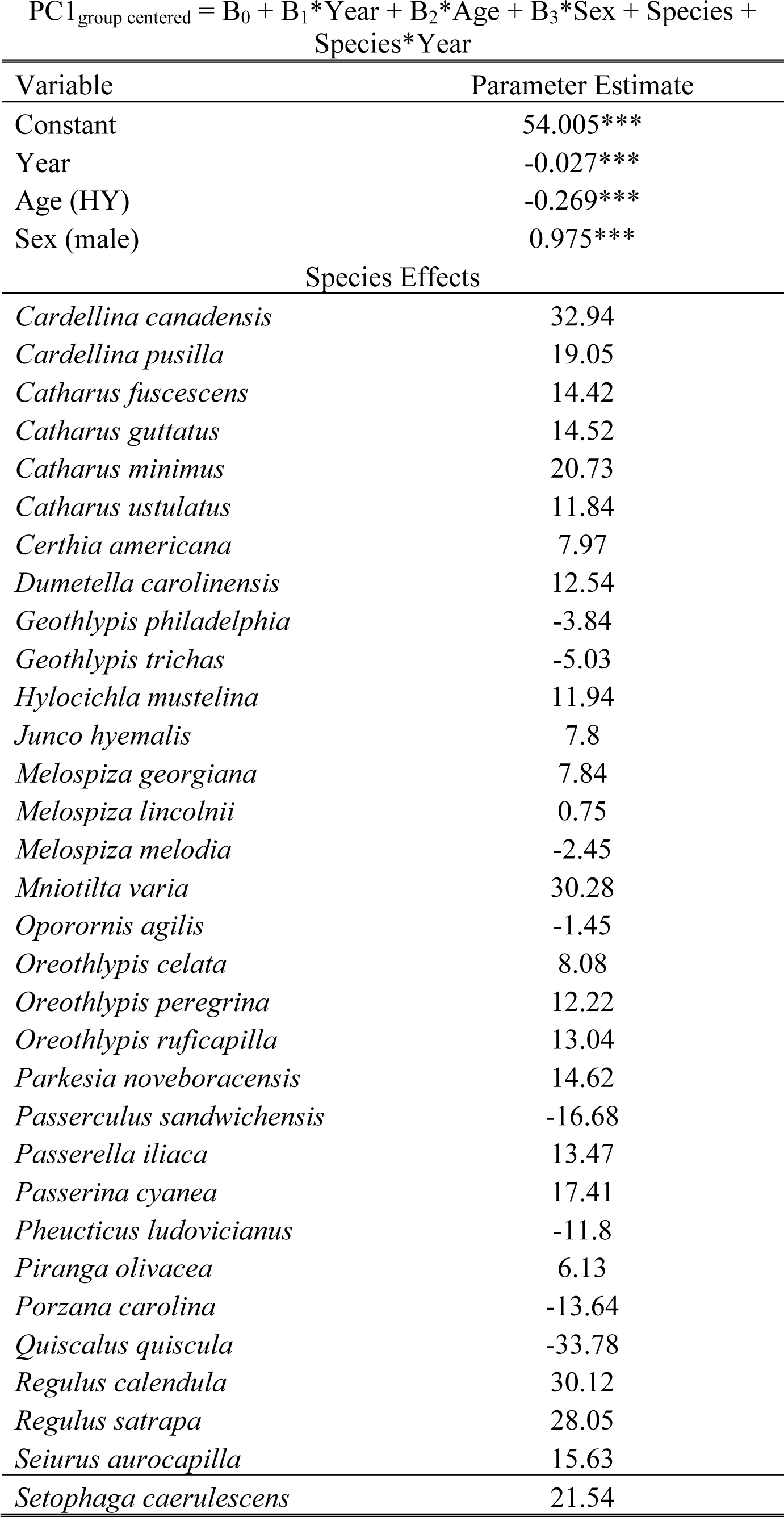
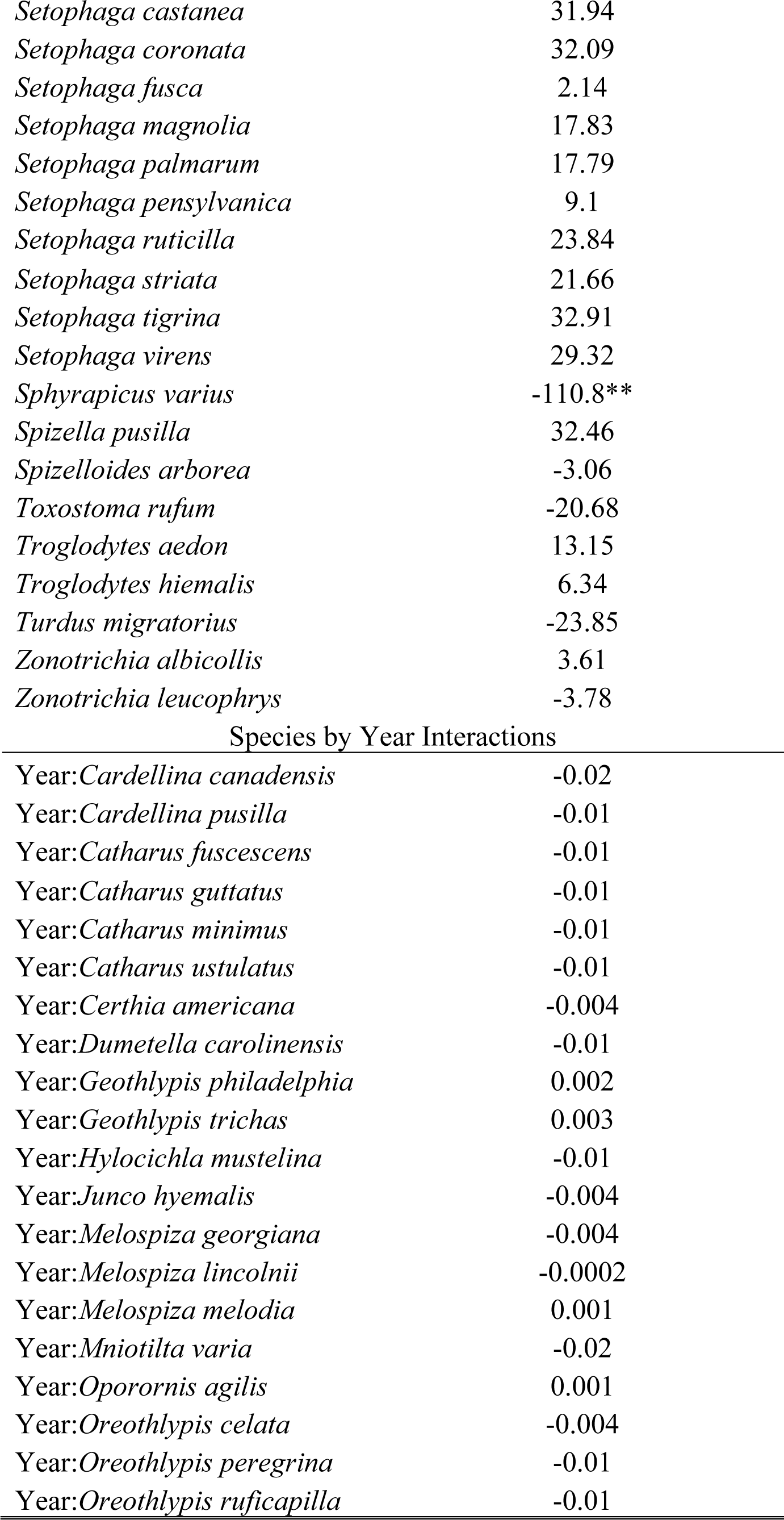
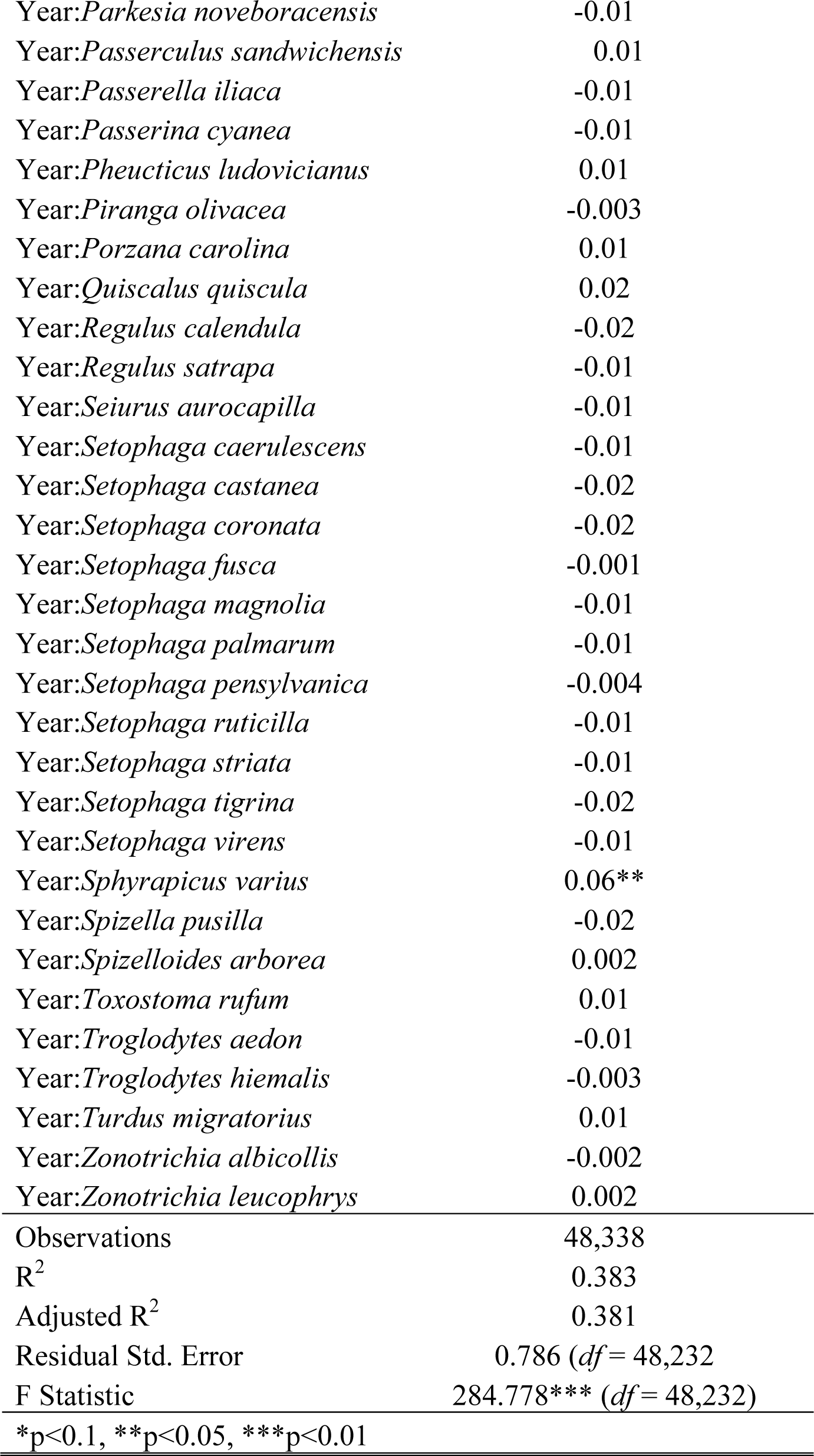
*PC1 Shows Decline in Body Size through Time*. All species and species by year interaction terms are relative to the reference taxon *Ammodramus savannorum*.

**Supplementary Table 5.**
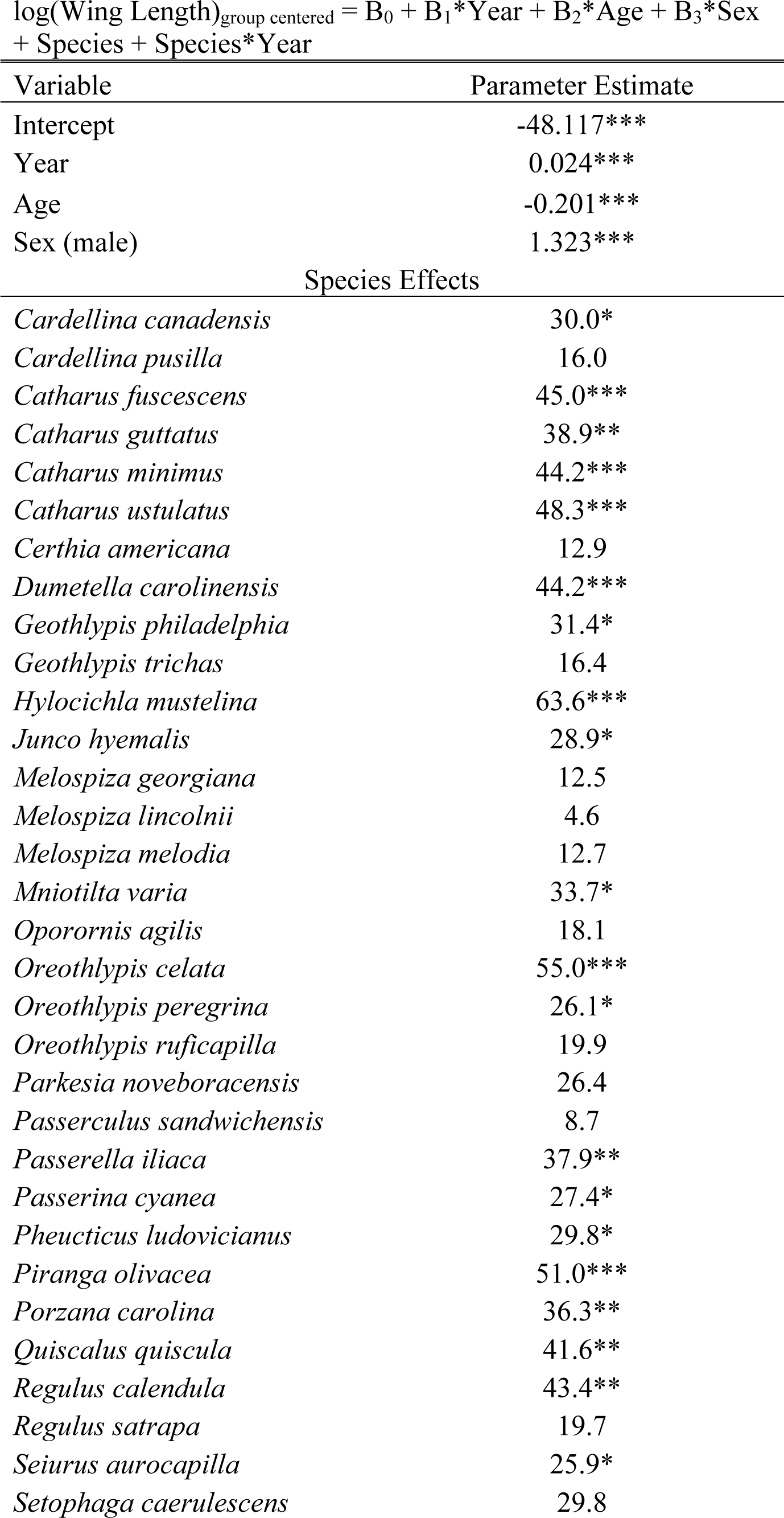
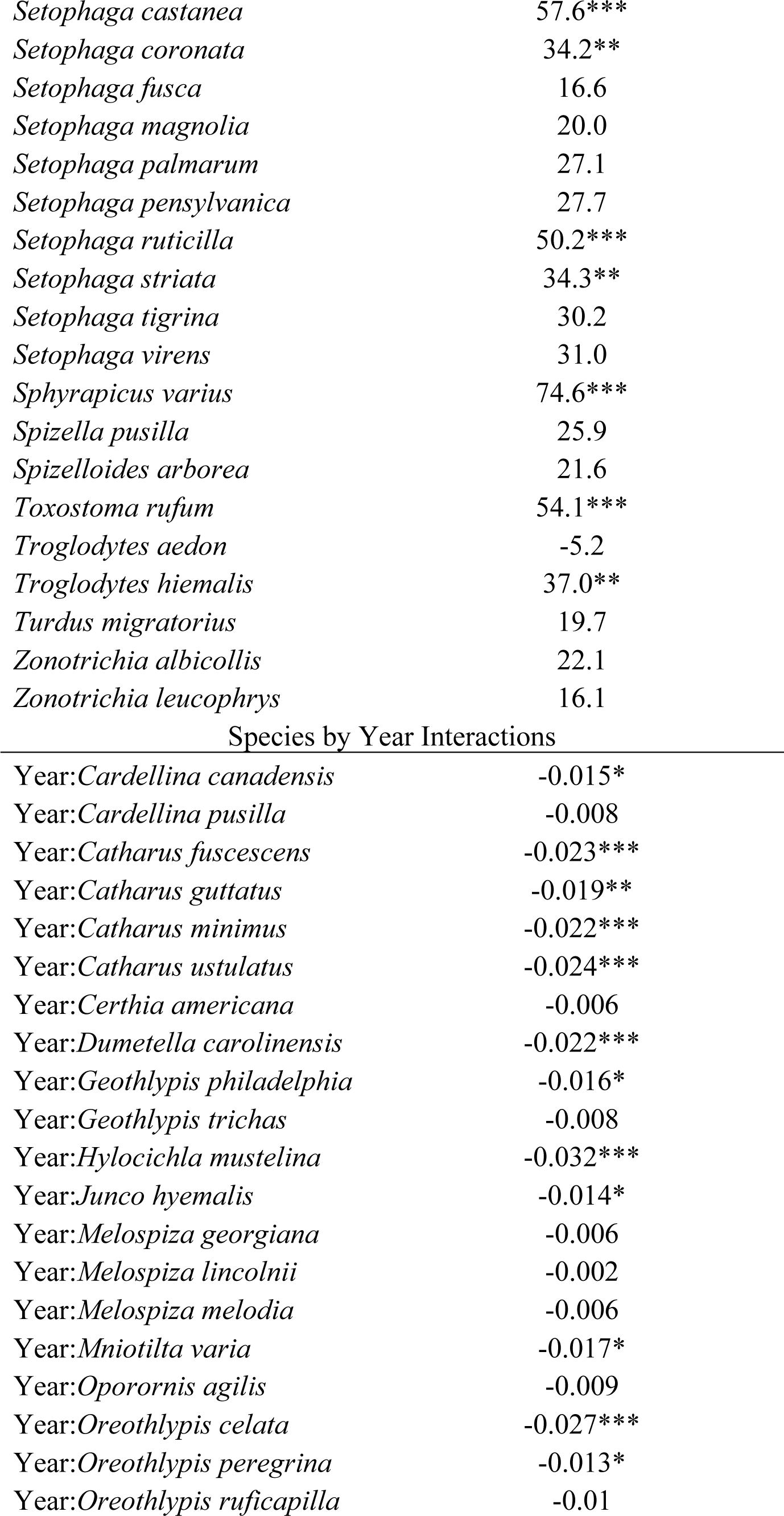
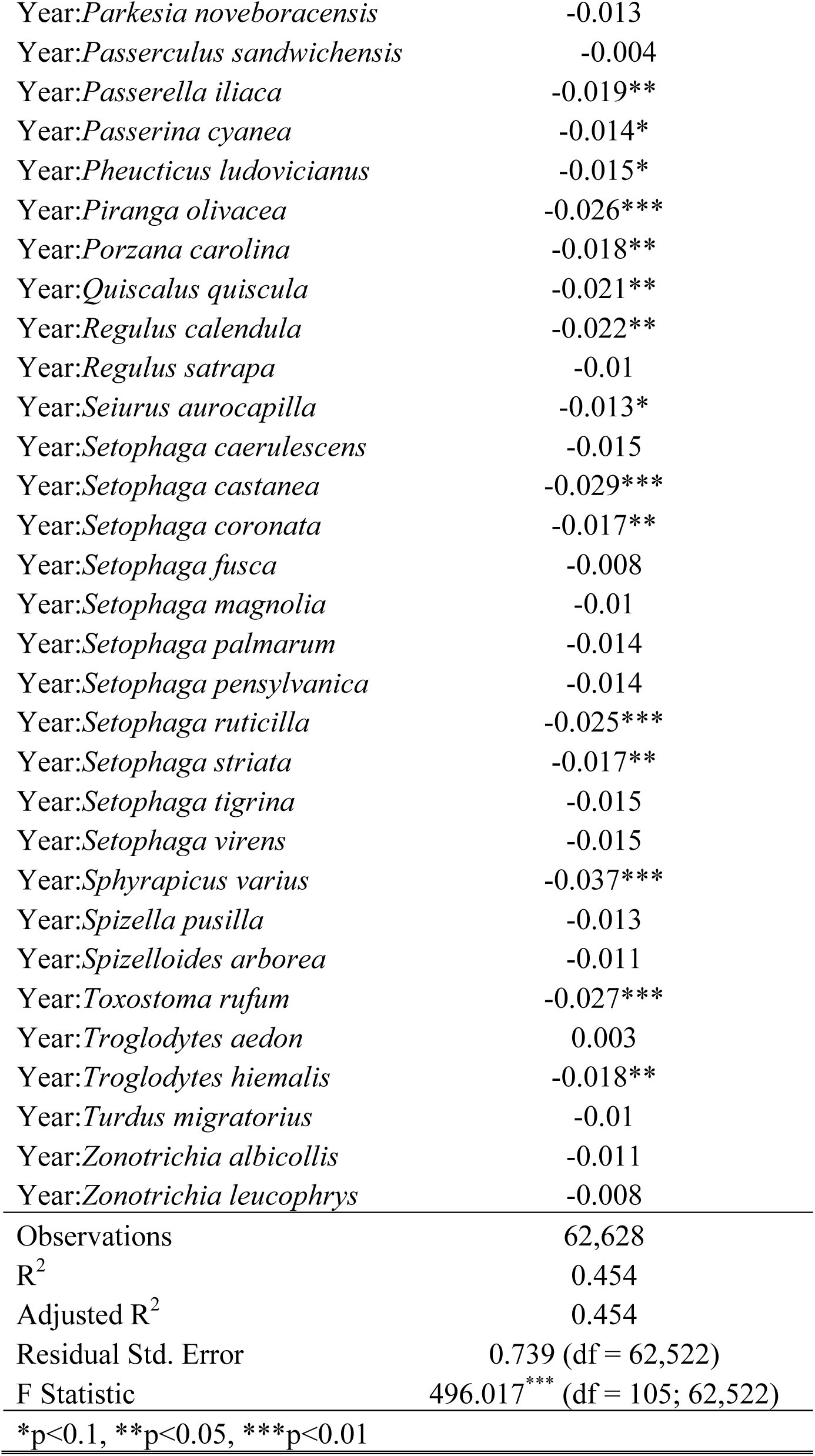
*Wing Length has Increased through Time*. All species and species by year interaction terms are relative to the reference taxon *Ammodramus savannorum*.

**Supplementary Table 6.**
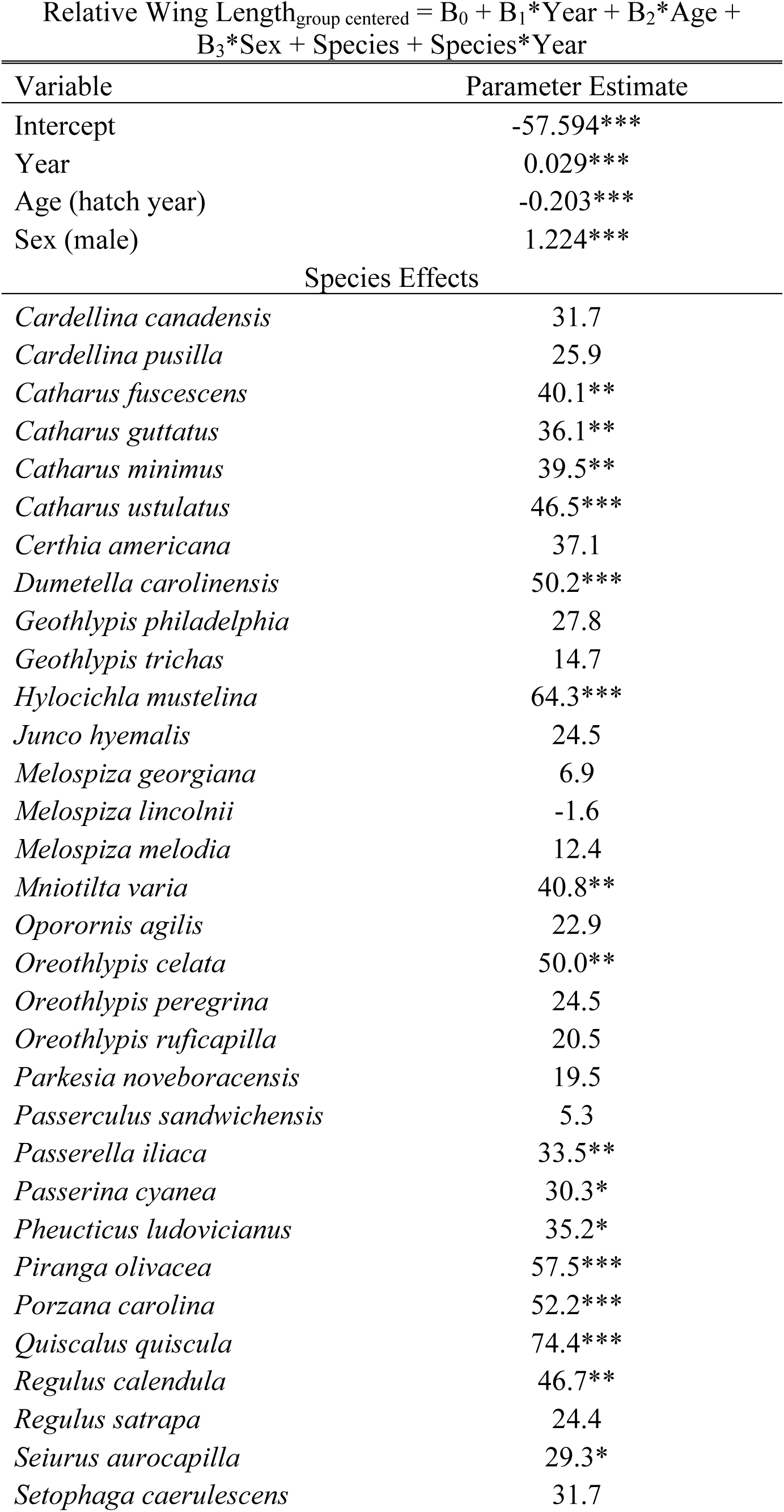
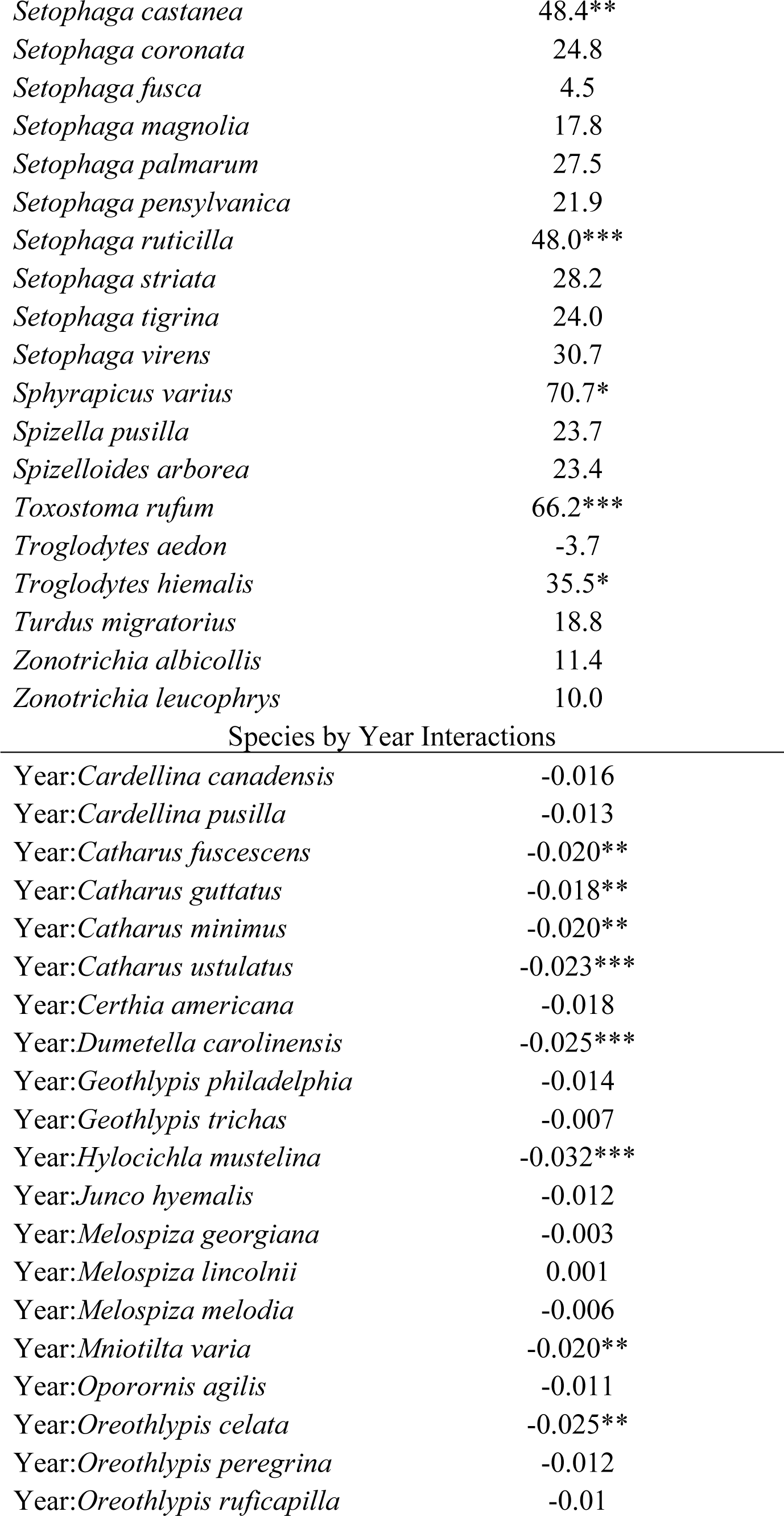
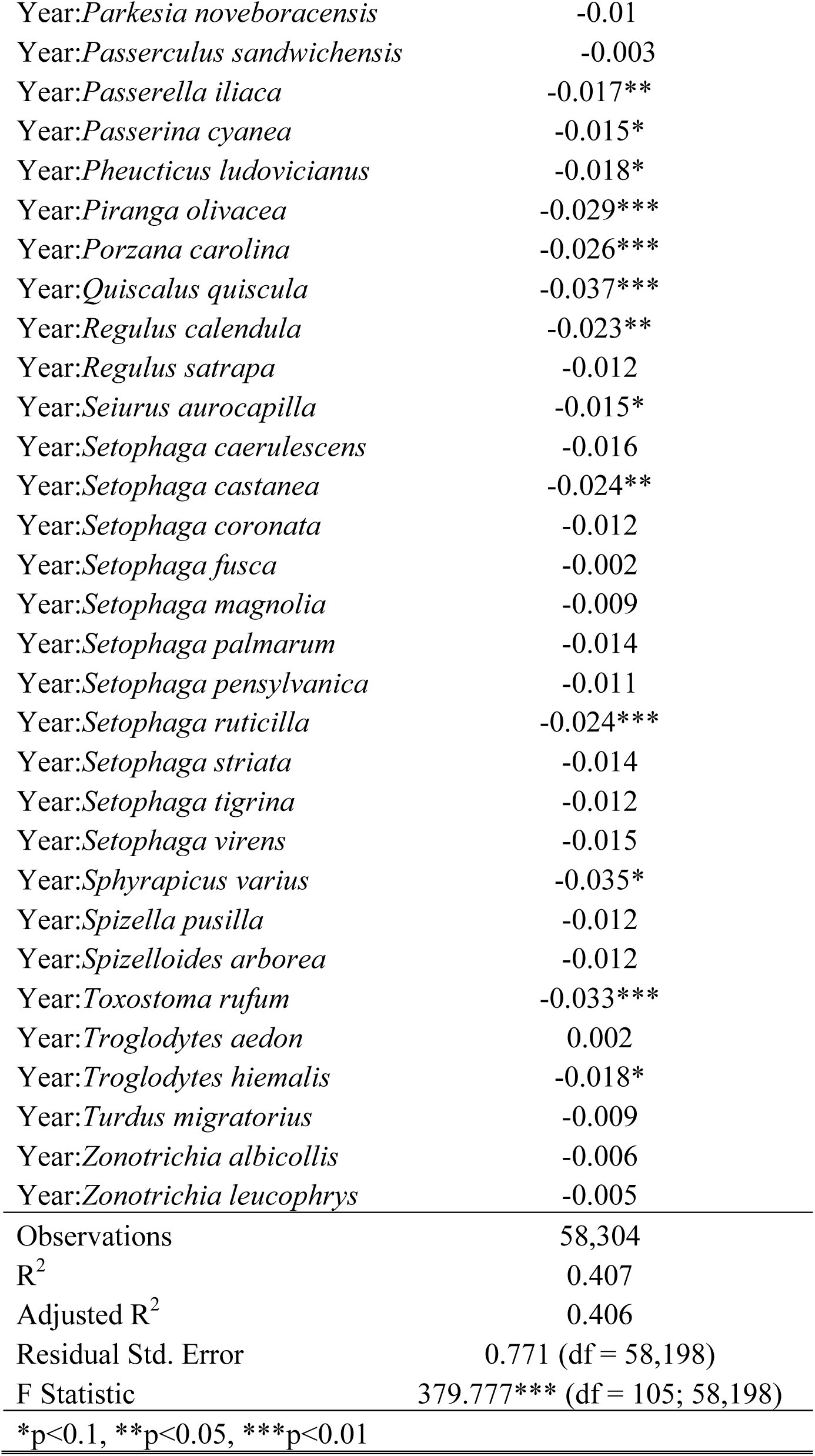
*Relative Wing Length has Increased through Time*. All species and species by year interaction terms are relative to the reference taxon *Ammodramus savannorum*.

**Supplementary Table 7.**
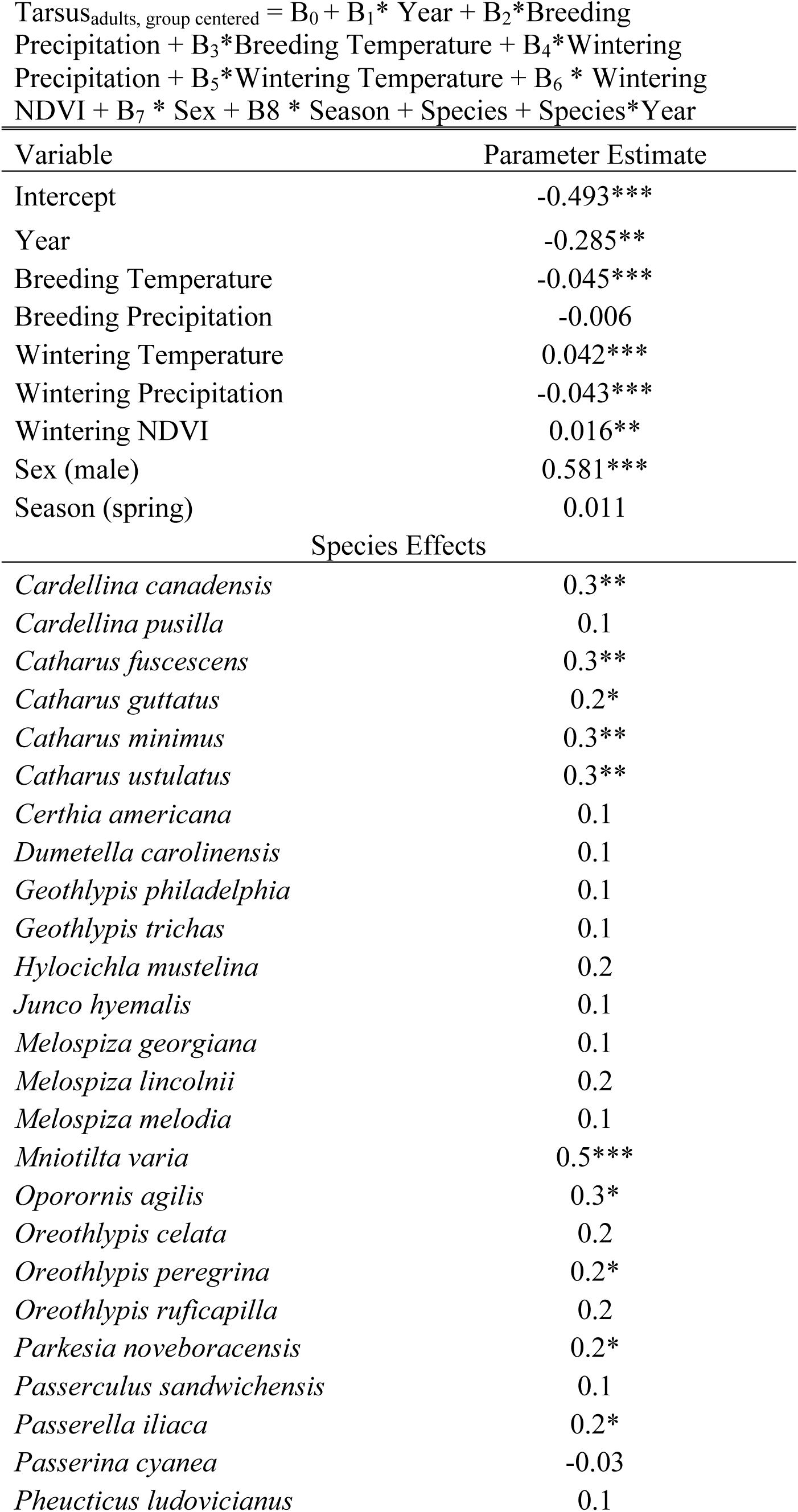
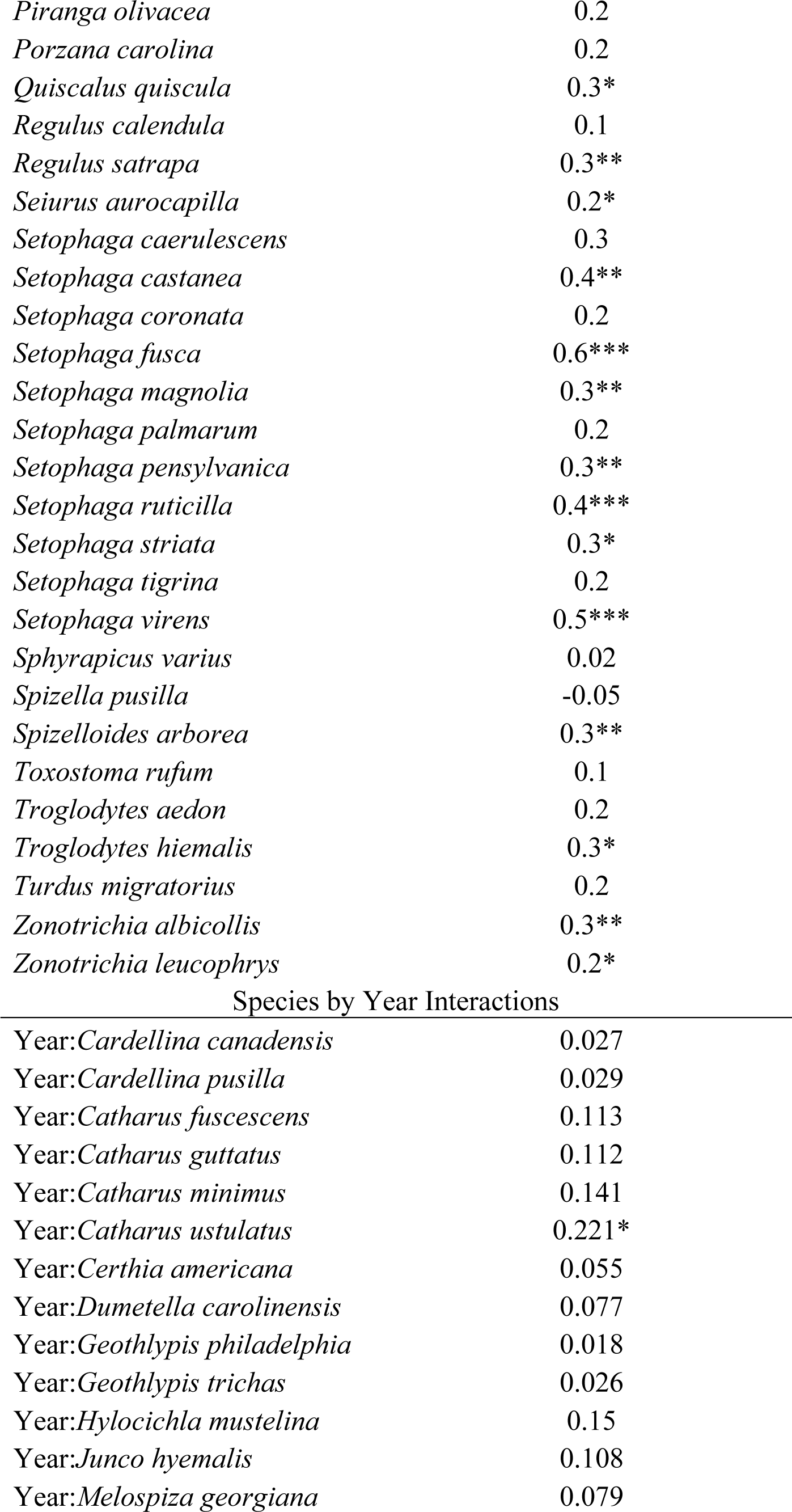
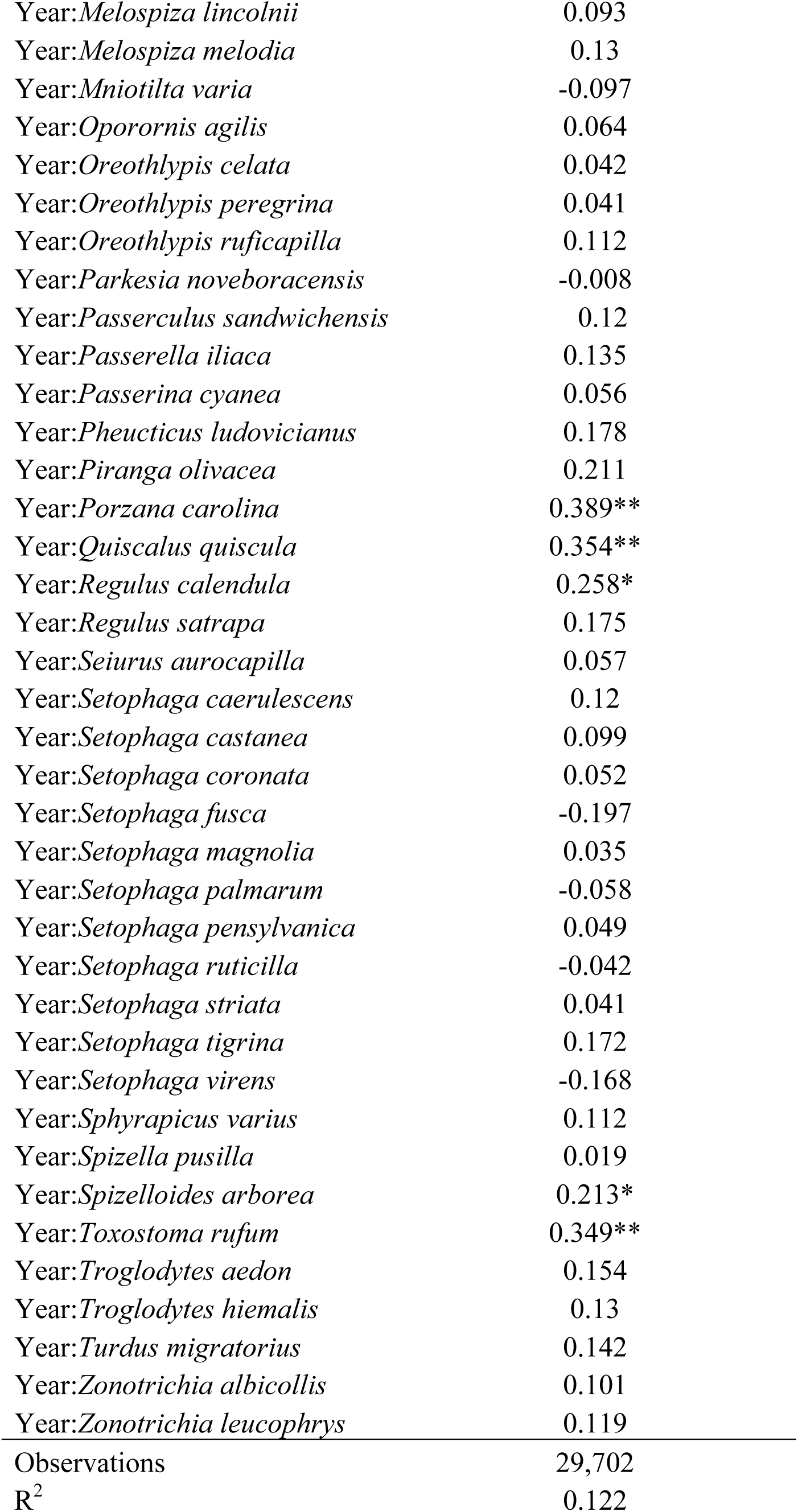
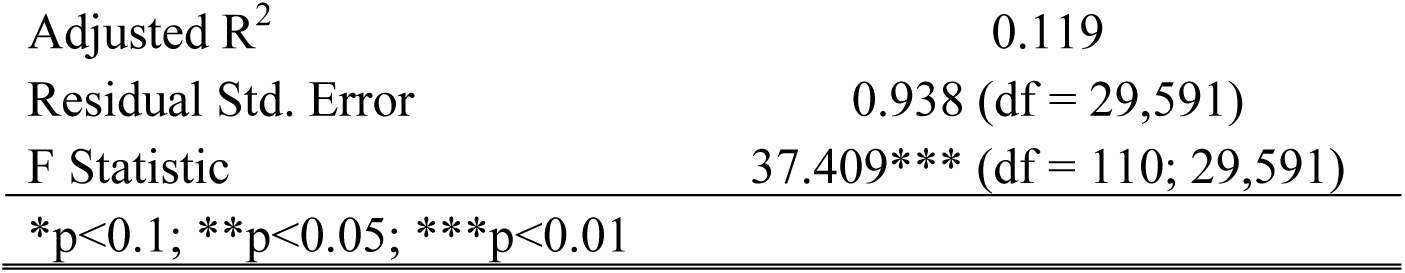
*Tarsus as a Function of Environmental and Climatic Variables on the Breeding and Wintering Grounds*. All species and species by year interaction terms are relative to the reference taxon *Ammodramus savannorum*.

